# ARP2/3 complex associates with peroxisomes to participate in pexophagy in plants

**DOI:** 10.1101/2022.04.07.487451

**Authors:** Jan Martinek, Petra Cifrová, Stanislav Vosolsobě, Jana Krtková, Lenka Sikorová, Kateřina Malínská, Zdeňka Mauerová, Ian Leaves, Imogen Sparkes, Kateřina Schwarzerová

## Abstract

ARP2/3 is a heteroheptameric protein complex evolutionary conserved in all eukaryotic organisms. Its conserved role is based on the induction of actin polymerization at the interface between membranes and the cytoplasm. Plant ARP2/3 has been reported to participate in actin reorganization at the plasma membrane during polarized growth of trichomes and at the plasma membrane-endoplasmic reticulum contact sites. We demonstrate here that individual plant subunits of ARP2/3 fused to fluorescent proteins form motile dot-like structures in the cytoplasm that are associated with plant peroxisomes. ARP2/3 dot structure is found at the peroxisome periphery and contains assembled ARP2/3 complex and WAVE/SCAR complex subunit NAP1. This dot occasionally colocalizes with the autophagosome, and under conditions that affect the autophagy, colocalization between ARP2/3 and the autophagosome increases. ARP2/3 subunits co-immunoprecipitate with ATG8f marker. Since mutants lacking functional ARP2/3 complex have more peroxisomes than WT, we link the ARP2/3 complex on peroxisomes to the process of peroxisome degradation by autophagy called pexophagy. Additionally, several other peroxisomal proteins colocalize with ARP2/3 dot on plant peroxisomes. Our results suggest a specific role of ARP2/3 and actin in the peroxisome periphery, presumably in membrane remodelling. We hypothesize that this role of ARP2/3 aids processes at the peroxisome periphery such as peroxisome degradation through autophagy or regulation of peroxisomal proteins localization or function.

**Significance statement:** ARP2/3 complex-positive dots associate exclusively with peroxisomes in plant cells, where it colocalizes with autophagosome marker ATG8f and several other proteins. Our experiments link ARP2/3 to pexophagy: colocalization between ARP2/3 dots and autophagosome increases when autophagy processes are induced or inhibited; ARP2/3 and ATG8f colocalize and co-immunoprecipitate, and finally, ARP2/3 mutants’ cells contain more peroxisomes than WT. Our results suggest a novel role of ARP2/3 in peroxisome structure and function regulation.

## INTRODUCTION

ARP2/3 complex is an evolutionary conserved heteroheptameric protein complex with a size of 220 kDa (Kollmar et al., 2012). The complex consists of two actin-related proteins ARP2 and ARP3, and five other subunits ARPC1-5 (actin-related protein complex components), not related to actin (Welch et al., 1997). Upon activation of the complex with nucleation-promoting factors (Rotty et al., 2013), the complex assists in the ATP-dependent nucleation of actin filaments (Welch et al., 1998). In plants, the ARP2/3 complex has been shown to be activated by the upstream regulatory pathway involving the SCAR/WAVE complex (for review see (Yanagisawa et al., 2013).

In animal cells, the ARP2/3 complex is essential for the formation of lamellipodia through the polymerization of actin filaments at the leading edge of the cell, and therefore for cell motility (Welch et al., 1997; Pollard and Borisy, 2003). This type of movement is necessary for cell migration during embryogenesis (Sawa et al., 2003), neuronal axon growth (Korobova and Svitkina, 2008; Kalil and Dent, 2014) or chemotaxis of white blood cells (Zicha et al., 1998). In yeasts, the ARP2/3 is involved in the formation of actin patches, dynamic actin structures in the cortical cytoplasm, important for endocytosis and cell wall remodeling (Young et al., 2004). In contrast to non-plant cells, plants lacking subunits of ARP2/3 complex are vital with specific defects in morphogenesis of cells with complex shapes, such as trichomes and epidermal pavement cells (for review see (Ivakov and Persson, 2013)).

Mutants lacking functional ARP2/3 complex have shorter hypocotyls (Mathur et al., 2003a; Kotchoni et al., 2009), shorter roots on sucrose-depleted medium (Dyachok et al., 2011), and cell-cell adhesion defects (Mathur et al., 2003a, 2003b; El-Din El-Assal et al., 2004). ARP2/3 mutants suffer from specific cell wall defects (Dyachok et al., 2008; Yanagisawa et al., 2015; Sahi et al., 2018) and dynamics of auxin transport (Sahi et al., 2018) (García-González et al., 2020). To understand the role of ARP2/3 in plants, it is essential to study the subcellular localization of the complex. The immunohistological approach to visualising the ARP2/3 complex in plant cells revealed the complex in close proximity to actin filaments (Fiserová et al., 2006; Maisch et al., 2009; Yanagisawa et al., 2015) or both actin and microtubules (Zhang et al., 2013a); (Havelková et al., 2015). ARP2/3 complex subunits and components of the SCAR/WAVE complex were observed to colocalize with various organelles such as the nucleus or ER (Zhang et al., 2013b). *In vivo* visualization of GFP-tagged ARPC5 subunit in *Arabidopsis thaliana* trichomes showed active ARP2/3 complex localized to the tips of growing trichome branches (Yanagisawa et al., 2015, 2018), which is consistent with its role in cell wall building. SCAR/WAVE subunit NAP1 was demonstrated to form puncta colocalizing with autophagosome marker ATG8, presumably regulating autophagy in plant cells (Wang et al., 2016). A study by Wang *et al*. ((Wang et al., 2019) further strengthened the link between ARP2/3 complex and autophagy in plants, showing that AtEH1/Pan1 protein recruited endocytic machinery components, clathrin, actin and ARP2/3 to autophagosomes at the PM. ARP2/3-nucleated actin assisting membrane remodelling during autophagy has been previously reported in yeast (Monastyrska et al., 2008) and mammalian cells (Kast et al., 2015; Coutts and La Thangue, 2016; Kast and Dominguez, 2015), suggesting that autophagy-related ARP2/3 functions are conserved. The above-mentioned localization studies suggest multiple roles of the ARP2/3 complex in plants.

We show here that ARPC2 and ARPC5 subunits of the ARP2/3 complex are localized in dot-like structures in the cytoplasm of tobacco and *Arabidopsis thaliana* cells, which are exclusively associated with peroxisomes. We further demonstrate that the peroxisome-associated dots constitute the whole assembled complex including a NAP1 subunit of activation complex SCAR/WAVE. Peroxisomes of ARP2/3 mutant *Arabidopsis* lines are larger and they are more abundant than in WT, which suggests a defect in peroxisome degradation. We tested this hypothesis and found out that peroxisomal ARP2/3 dots colocalized with autophagosomal marker ATG8f, and that the formation of ARP2/3 dots was sensitive to autophagy induction and inhibition. Since ARP2/3 subunits co-immunoprecipitated with the ATG8f marker, we concluded that the ARP2/3 complex is involved in the pexophagy of plant peroxisomes. Interestingly, several other peroxisomal proteins colocalized to the ARP2/3-positive peroxisomal subcompartment. Our results open a possibility that the ARP2/3-nucleated actin cytoskeleton assists multiple peroxisome functions at the peroxisomal periphery.

## RESULTS

### ARP2/3 complex forms motile dots associated with peroxisomes

We prepared ARP2/3 complex subunits GFP-*At*ARPC2 and GFP-*At*ARPC5 and expressed them under 35S promoter, and we used GFP-*Nt*ARPC2 in the β-estradiol inducible system (XVE::GFP-*Nt*ARPC2, (Havelková et al., 2015). GFP-ARPC2 and GFP-ARPC5 subunits of ARP2/3 complex localized into small motile dots, and a diffuse cytoplasmic signal (Figure 1a-c) in *Arabidopsis* cells. We used various organelles markers to test if ARP2/3 subunits-positive dots associate with a specific organelle (Figure S1). We show here that ARP2/3 subunits are exclusively colocalized with peroxisomes marked by mCherry-PTS1 (Figure 1d-f). We found out that all three tested fusion proteins colocalized with peroxisomes in stably transformed *Arabidopsis thaliana* (Figure 1d-f) and BY-2 tobacco cultured cells (Figure S2e), as well as in infiltrated *Nicotiana benthamiana* leaves (Figure S2f). The dots were always associated with peroxisomes and no free ARP2/3 subunits-positive dots were noted. The analysis of epidermal cells in hypocotyl showed that 73.7% (SD = 16,2%) of peroxisomes were associated with ARP2/3, with relatively high observed variability among various experiments. Further, peroxisomes were associated with only one single ARP2/3 dot; more dots associated with one peroxisome we observed only scarcely.

**figure 1.**
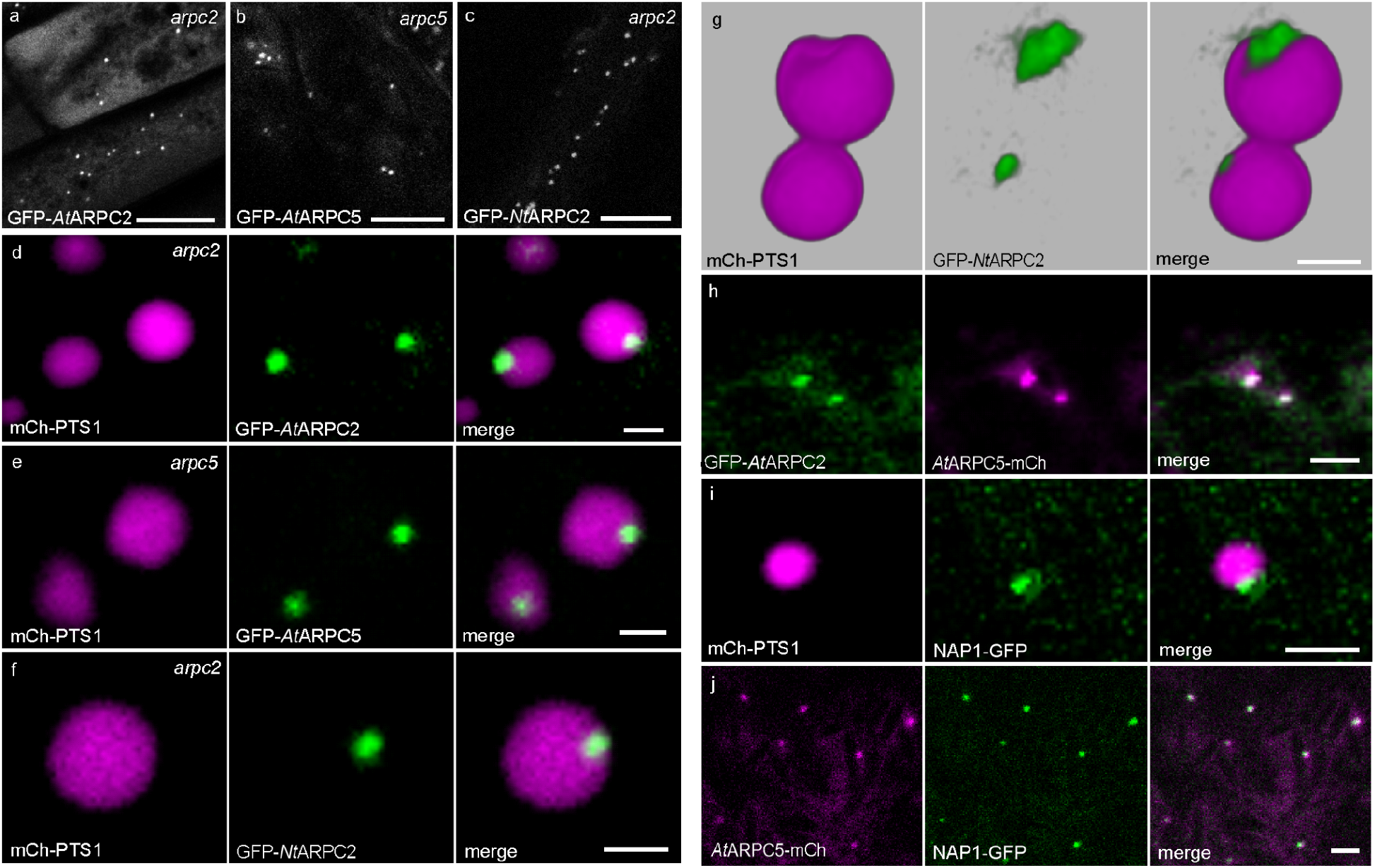
ARP2/3 complex subunits GFP-ARPC2 and GFP-ARPC5 and NAP1-GFP, a subunit of an ARP2/3 activating complex, colocalize with peroxisomes. (a) GFP-*At*ARPC2, (b) GFP-*At*ARPC5 and (c) inducible GFP-*Nt*ARPC2 form small motile dots in hypocotyl cells of transgenic *Arabidopsis thaliana*. (d) GFP-*At*ARPC2, (e) GFP-*At*ARPC5 and (f) GFP-*Nt*ARPC2 motile dots colocalize exclusively with peroxisomes, visualized by the mCherry-PTS1 marker in *Arabidopsis thaliana* hypocotyl epidermis cells. (g) Superresolution Airyscan imaging of ARP2/3 subunit GFP-*Nt*ARPC2 (green) and peroxisome (magenta) -3D view. (h) GFP-*At*ARPC2 and *At*ARPC5-mCherry colocalize in motile dots, suggesting that peroxisome-associated dots contain assembled ARP2/3 complexes. (i) NAP1, a subunit of ARP2/3 activating complex SCAR/WAVE, localizes to motile dots associated with peroxisomes in *Arabidopsis* stably expressing cells as well. (j) ARPC5-mCherry and NAP1-GFP colocalize to motile dots, suggesting that peroxisome-associated dots contain activated ARP2/3 complexes. Scale bars: a - c = 10 µm, d - g = 1 µm, h - j = 2 µm.

Motile dots of inducible GFP-*Nt*ARPC2 were observed also in cells induced by very low concentrations of β-estradiol (Figure S2a-d), which suggests that the dots are not overexpression artifacts. GFP-*At*ARPC2 and GFP-*At*ARPC5 expressed in respective *arpc2* and *arpc5* mutant backgrounds complemented the phenotype of distorted trichomes, indicating that the GFP-tagged subunits were fully functional (Figure S2g-k). The complementation of the *arpc2* phenotype by the inducible vector carrying GFP-*Nt*ARPC2 has been demonstrated previously (Havelková et al., 2015).

ARP2/3-positive dots were always associated with the periphery of the peroxisome. The peripheral localization to the peroxisome is especially evident when ARP2/3-dots and peroxisomes move. While the peroxisome was transported through the cytoplasm, the ARP2/3 dot was turning around the peroxisome (Supporting movie S1). The dot seemed to be closely associated with the outer surface of the peroxisome (Figure 1d-f), but also a more loose association of the dot with the peroxisome was rarely observed (Figure S2e). We analyzed the colocalization of ARP2/3 dot with peroxisomes using Airyscan superresolution microscopy. Analysis showed peripheral localization of the ARP2/3 structure at the peroxisome (Figure 1g). The ARP2/3-positive structure that associated with the peroxisome periphery often deformed the mCherry-PTS1-labeled compartment, giving the impression of pushing on the peroxisomal periphery or membrane (Figure 1g).

### Motile dots associated with peroxisomes contain activated ARP2/3 complex and peroxisomal ARP2/3 co-sediments with peroxisomal proteins

Since ARPC2 and ARPC5 are components of the same protein complex, they should colocalize in the ARP2/3-positive peroxisome-associated dot. We thus co-expressed *At*ARPC5-mCherry fusion protein with GFP-*At*ARPC2 in *Arabidopsis*. Both subunits colocalized to the same spot (Figure 1h). To test if ARP2/3 positive structures associated with peroxisomes contain activated ARP2/3 complex, we co-expressed *Arabidopsis* NAP1, a subunit of plant WAVE/SCAR complex, with a peroxisomal marker. NAP1-GFP motile dots colocalized with peroxisomes as a small motile dot, similarly to ARPC5 and ARPC2 (Figure 1i). Indeed, both markers colocalized in cells expressing NAP1-GFP with *At*ARPC5-mCherry to the same motile dots (Figure 1j). To confirm microscopic observations, we performed co-immunoprecipitation (co-IP) experiments, where anti-GFP antibody-labeled magnetic beads were used for immunoprecipitation of GFP-*At*ARPC2 or NAP1-GFP in protein extracts from plants co-expressing *At*ARPC5-mCherry. *At*ARPC5-mCherry co-immunoprecipitated with GFP-*At*ARPC2 and NAP1-GFP (Figure S2l). Since the two ARP2/3 subunits co-immunoprecipitated with each other, and the ARPC5 subunit co-immunoprecipitated with NAP1, we assumed that peroxisomes associated dots consist of the assembled and activated complex.

To understand the role of ARP2/3 complex localization on peroxisomes, we aimed to identify protein interactors of the ARP2/3 complex in the ARP2/3 dots, associated with peroxisomes. We prepared protoplasts from BY-2 cells expressing β-estradiol inducible GFP-*Nt*ARPC2, which we used for the preparation of peroxisome-enriched fraction using centrifugation on a sucrose gradient. The peroxisome-enriched fraction was used for the GFP-co-IP and isolated proteins were analyzed using nLC-MS^2^. Free GFP expressing BY-2 cells and non-induced GFP-*Nt*ARPC2 cells were used as controls. This approach allowed us to identify 19 proteins likely interacting with GFP-*Nt*ARPC2 (Supporting Table 1). All ARP2/3 complex subunits were identified as specific interactors of ARPC2, which further confirms that the peroxisome-associated ARP2/3 dot is formed of assembled ARP2/3 complex. We also identified other proteins in the GFP-co-IP experiment (Supporting Table 1), which contained the PTS1 or PTS2 targeting sequence and thus represented peroxisome-targeted enzymes responsible for peroxisomal metabolism. For instance, several isoforms of peroxisomal fatty acid β-oxidation multifunctional protein AIM1-like and 4-coumarate--CoA ligase-like, enzymes responsible for β-oxidation of fatty acids and biosynthesis of jasmonates (Supporting Table 1) were identified in our analysis.

### Interaction of ARP2/3 dots and actin

Small, discrete, dot-like structures formed by ARP2/3 subunits were highly motile and moved along the actin cytoskeleton, but they often lacked a direct interaction with the actin bundle (Figure 2a, Supporting Movie S2). Such dynamics suggested that the ARP2/3 dot is not directly associated with actin bundles labeled with lifeact-mRFP (Figure 2a) or GFP-FABD (Supporting Movie S2). Colocalization with peroxisomal marker revealed that ARP2/3-positive dots were located on the surface of peroxisomes and moved together with them (Supporting Movie S1). During the peroxisome transport along actin filaments, the changing position of the associated ARP2/3 dot sparsely corresponded to the adjacent bundle of actin filaments in plants co-expressing GFP-*Nt*ARPC2, mCh-PTS1, and GFP-FABD (Supporting Movie S2). This explained the loose and occasional association of ARP2/3 with actin *in vivo*, where ARP2/3 dots were carried on peroxisomes as a cargo independent of actin filaments. We tested if ARP2/3 plays a role in peroxisome motility along actin filaments. The speed of peroxisomes measured in plants lacking functional ARP2/3 complex showed that the speed is comparable in WT and all tested mutants (Figure 2b), which suggests that the role of ARP2/3 on peroxisome does not plays a role in peroxisome motility. This was further confirmed in experiments with latrunculin B. After treatment of hypocotyl epidermal cells with 10 µM latrunculin B for 3 h, the motility of peroxisomes ceased, and the organelles tended to cluster (Figure 2c). However, ARP2/3 dots remained associated with peroxisomes and showed no structural or localization change, even though actin was completely depolymerized, as shown in plants expressing the GFP-FABD marker (Figure S3a, b). Similarly, the treatment with 40 µM oryzalin for 3 h had also no effect on ARP2/3 dot structure and localization (Figure S3c-e).

**figure 2.**
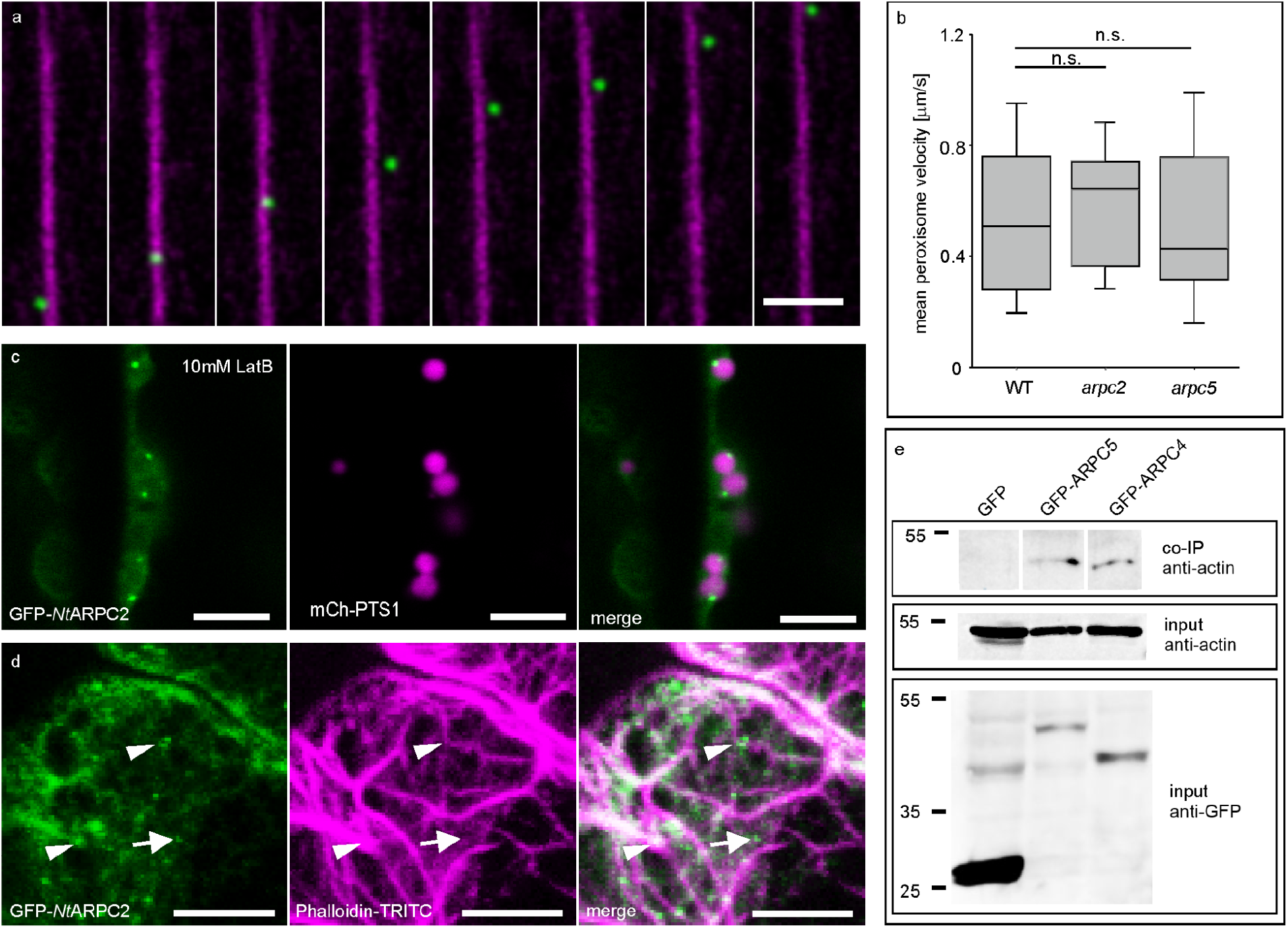
(a) ARP2/3 motile dot moves along actin filaments with occasional detachment from actin filaments (time step = 2.58 s per frame).(b) treatment of *Arabidopsis* cotyledon epidermal cells with 10 µM latrunculin B for 3 hours caused peroxisomes clustering, but had no effect on ARP2/3 dot localization on the peroxisomes. (c) Visualisation of actin filaments in fixed BY-2 cells using TRITC-phalloidin. In BY-2 cells, ARP2/3 dots represented by GFP-*Nt*ARPC2 colocalize with thick actin filaments (arrowheads) or diffuse actin signal (arrow). **(d)** The speed of peroxisomal movement is not changed in *arpc2* and *arpc5* mutant lines compared to WT. The central line in boxplots represents the median; number of measurements (n) = 18 (WT), 14 (*arpc2*), 16 (*arpc5*). Each measurement represents the average velocity of all peroxisomes in one video, where tens of peroxisomes in several cells were captured. T-tests comparing WT and each mutant were not significant (*p* > 0.05). (**e**) Actin co-immunoprecipitates with GFP-ARPC5 and GFP-ARPC4 in total protein extracts from BY-2 stably expressing GFP-tagged proteins. Free GFP-expressing cells are used as a negative control. Scale bars: a = 3 µm, b, d = 5 µm.

Further, we tested the possibility that ARP2/3 dots are associated with finer actin filaments, which were not detectable using FABD-or Lifeact-based actin markers. We fixed BY-2 cells stably expressing GFP-*Nt*ARPC2 and visualized actin cytoskeleton using TRITC-labeled phalloidin. We found out that most ARP2/3 dots colocalized with actin filaments, or the dots were found in the close vicinity of actin filaments (Figure 2d). This is consistent with our observations of living cells, where peroxisomes with ARP2/3 dots were transported along with the actin cables. In fixed cells, we also observed dots, which were obviously not associated with thick actin bundles (Figure 2d, arrow). However, these dots were always colocalizing with a thin actin filament or were surrounded by a cloud of signal, representing presumably polymerized short or thin actin filaments. Considering the small size of ARP2/3 dots associated with peroxisomes and a very fine structure of short polymerized actin, the fluorescence microscopy method has a limited resolution in this case. We have used a co-IP experiment to test if our tagged subunits interact with actin. We were able to pull-down actin in total protein extracts from BY-2 cells expressing GFP-*At*ARPC4 and GFP-*At*ARPC5 proteins (Figure 2e). Our results suggest that GFP-tagged subunits of ARP2/3 interact with actin, but the ARP2/3 function is not related to transport along actin filaments.

### Peroxisome-associated ARP2/3 dot structural integrity requires ARPC2 subunit

We investigated if the ARP2/3 dot structure requires the presence of all ARP2/3 subunits. We expressed GFP-ARPC2 in *Arabidopsis arpc5* mutant and GFP-ARPC5 in the *arpc2* mutant and compared ARP2/3 localization. The signal of GFP-ARPC2 in the *arpc5* mutant background was detected as a discrete, peroxisome-associated dot (Figure 3a). Interestingly, GFP-ARPC5 in the *arpc2* mutant background frequently formed discrete dots, but the signal was also dispersed in the peroxisome (Figure 3b). This suggests that the presence of the ARPC2 subunit is needed for the targeting of ARP2/3 subunits to a peripheral dot associated with peroxisomes. This is also consistent with the fact that the ARPC2 subunit forms, along with the ARPC4 subunit, the core of the complex (Robinson et al., 2001).

**figure 3.**
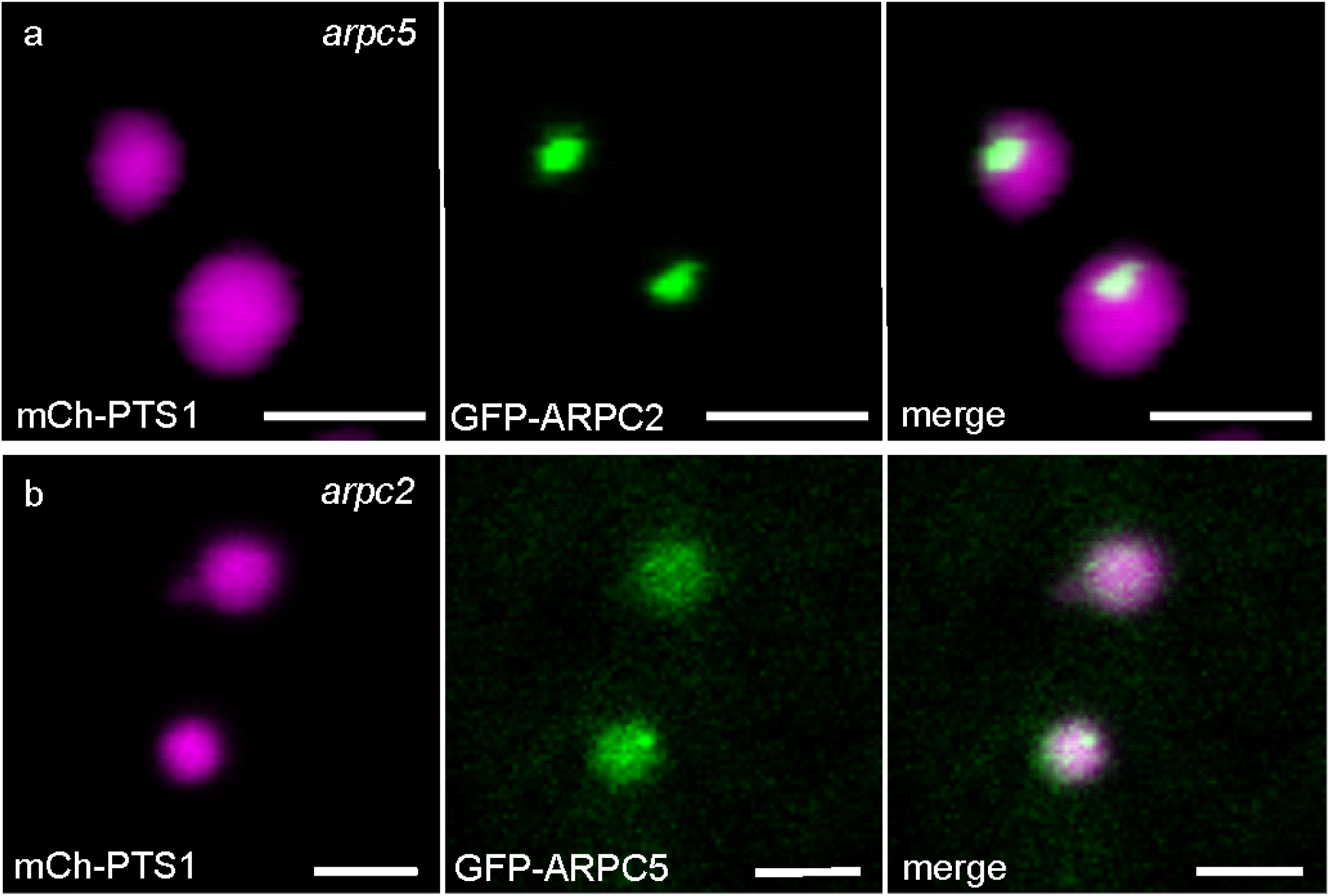
GFP-ARPC5 localization pattern in peroxisomes is dependent on the presence of the ARPC2 subunit.(a) GFP-ARPC2 forms discrete dots associated with peroxisomes in the *arpc5* mutant background. (b) GFP-ARPC5 signal in *arpc2* mutant background still forms dots associated with peroxisomes, but the signal is also dispersed. Scale bars = 2 μm.

### Peroxisomes in mutants lacking ARP2/3 complex are more abundant and larger, but metabolically functional

Since peroxisome motility was unchanged in ARP2/3 mutants, we searched for the structural or functional differences of peroxisomes in WT and ARP2/3 mutant *Arabidopsis* plants. Using the peroxisomal marker mCherry-PTS1 we found out that peroxisomes are significantly more abundant in *arpc2* and *arpc5* mutants (Figure 4a). Further analysis showed that the peroxisomes in *arpc2* and *arpc5* mutants were significantly larger than in WT (Figure 4b, c, respectively). We further tested if peroxisomal functions are impaired in ARP2/3 mutants lacking functional ARP2/3 complex. Peroxisomes are the site of β-oxidation in plants, and one of the methods used for testing this function is the effect of indole-3-butyric acid (IBA). IBA is ineffective in plant cells, but when converted to auxin (IAA) through the β-oxidation pathway located in peroxisomes, it affects the growth of seedlings (Adham et al., 2005). Plants impaired in β-oxidation are resistant to IBA because they do not properly convert IBA to auxin. ARP2/3 mutants grown on a medium with IBA showed clear growth inhibition (Figure 4d) and increased initiation of lateral roots (Figure S4a), which suggests that they are able to sufficiently convert IBA to auxin. Peroxisomes are also involved in the mobilization of storage compounds during *Arabidopsis* seeds germination, and mutants impaired in these functions are not able to germinate on media without sucrose (Graham, 2008). Since ARP2/3 mutants germinate normally on media without sucrose and oil bodies degradation during germination in dark without sucrose was not affected (Figure S4b, c), these peroxisomal functions are not impaired in ARP2/3 mutants. Therefore, peroxisomes in ARP2/3 mutants show structural changes, but they are still competent to perform β-oxidation.

**figure 4.**
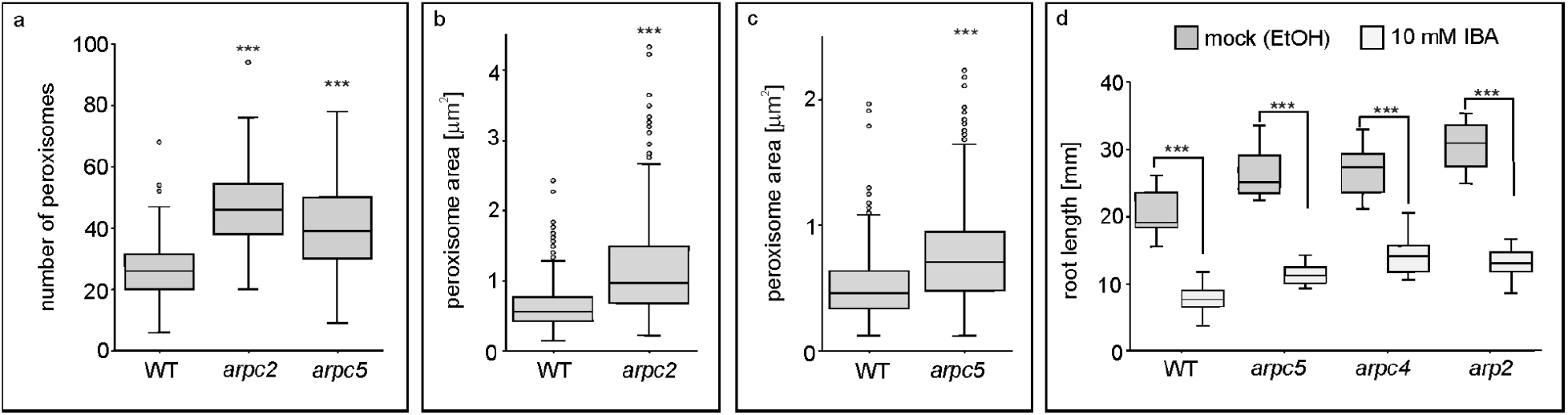
Cells of ARP2/3 mutants have more abundant, larger and functional peroxisomes. **(a)** *arpc2* and *arpc5* mutants contain more peroxisomes compared to WT. The number of analyzed pictures (n) = 129 (WT), 32 (*arpc2*), 47 (*arpc5*). **(b, c)** Peroxisomes in both *arpc2* and *arpc5* mutants are significantly larger than in WT plants. The number of measured peroxisomes (n) for graph b is 282 (WT) and 386 (*arpc2*), 10 plants in each variant were analyzed. For graph c, n = 521 (WT) and 447 (*arpc5*). (d) ARP2/3 mutant lines a*rp2, arpc4* and *arpc5* respond to IBA by reducing root growth, indicating a functional β-oxidation pathway. Number of analyzed roots (n) = 17 (WT mock), 18 (WT IBA), 9 (*arpc5* mock), 8 (*arpc5* IBA), 10 (*arpc4* mock), 9 (*arpc4* IBA), 10 (*arp2* mock), 10 (*arp2* IBA). Central lines in boxplots represent the median, the statistical significance of T-tests is indicated by asterisks (*p* < 0.001 = ***).

### Peroxisome-associated ARP2/3 dots colocalize with autophagosomes with low frequency

Since ARP2/3 has been previously shown to participate in autophagosome formation (Wang et al., 2016, 2019), our result may imply that the ARP2/3 complex is involved in the process of peroxisomes degradation, the pexophagy (Farmer et al., 2013). We co-expressed the autophagosomal marker ATG8f-RFP with GFP-*At*ARPC2 and found out that a subset of both markers colocalized (Figure 5a). We, therefore, generated a triple marker line, where CFP-PTS1 was co-expressed to visualize peroxisomes (Figures 5b and c). If peroxisome with both markers (ARP2/3 and ATG8f) was recorded, the two markers always colocalized in the same compartment (Figure 5c). Surprisingly, quantification analysis showed that ARP2/3 dots and autophagosomes colocalized on the same peroxisome with very low frequency. On average, 4.18 % of ARP2/3 dots colocalized with autophagosomes (n = 147 of analyzed optical fields). Video of plant cells expressing GFP-*At*ARPC2 and ATG8f-RFP revealed that the association of ARP2/3 dots and autophagosomes was usually transient, as ARP2/3 dots and autophagosomes were repeatedly attaching and detaching (Supporting Movie S3). The autophagosomes we observed in peripheral association with peroxisomes, colocalizing with ARP2/3 dots, were significantly smaller than the peroxisomes they were attached to (Figure 5c). Much less often we observed the engulfment of the whole peroxisome in the ATG8f-mRFP-marked membrane, suggesting an advanced step of pexophagy (Figure S5).

**figure 5.**
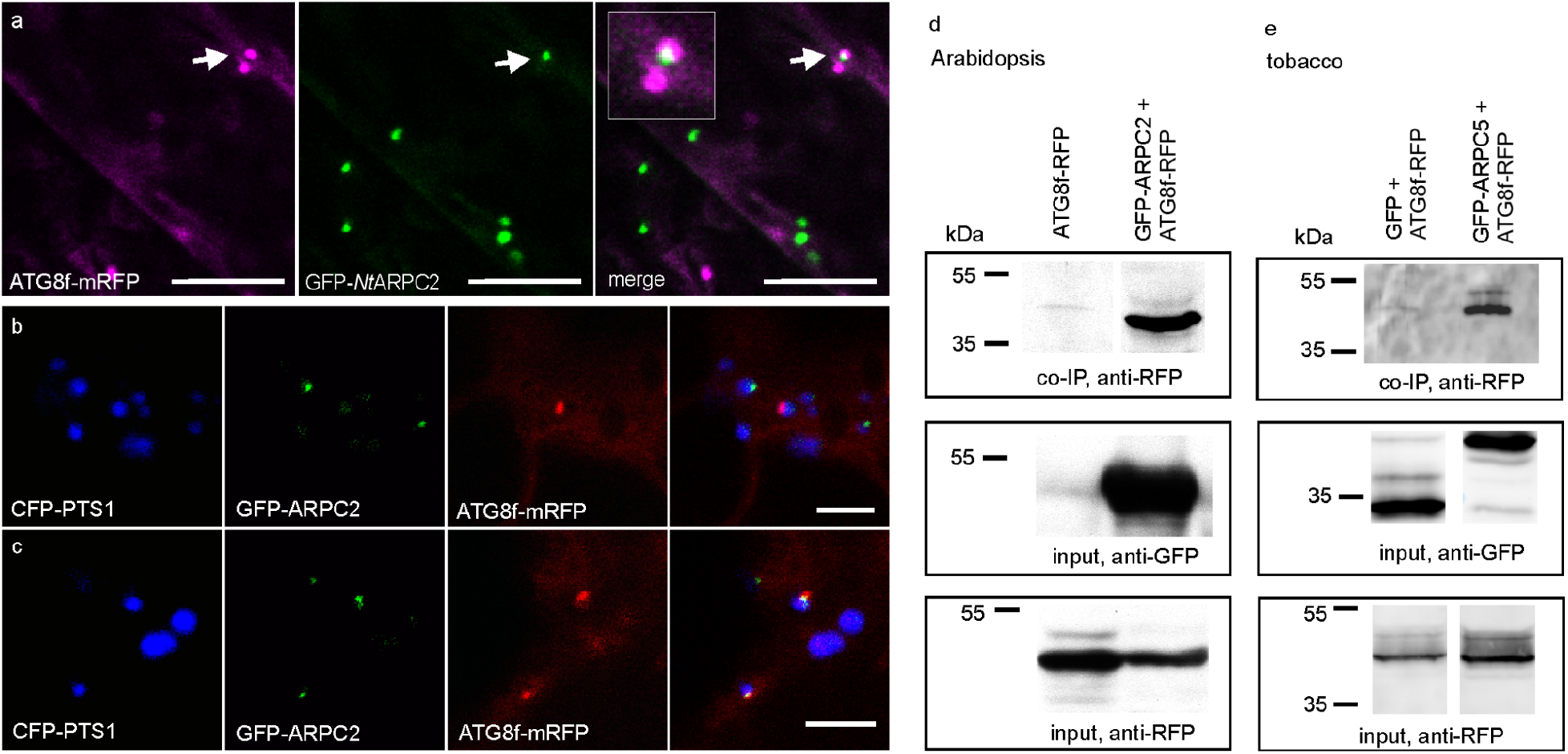
ARP2/3 colocalizes and interacts with autophagosome marker ATG8f and peroxisomes.(a) A representative picture of the ATG8f-mRFP and GFP-*Nt*ARPC2 double marker line shows that the two markers colocalize in the same compartment if found on the same peroxisome. (b) A representative picture showing colocalization of peroxisomal marker CFP-PTS1, ARP2/3 marker GFP-ARPC2 and ATG8fmRFP in the epidermis of triple marker line of *Arabidopsis* in hypocotyls. (c) ARP2/3 and ATG8f-mRFP colocalize to the identical compartment in peroxisomes in the triple marker line, if found on the same peroxisome. (d, e) ATG8f-mRFP co-immunoprecipitates with GFP-ARPC2 in *Arabidopsis* extracts expressing stably both markers, and with GFP-ARPC5 in transiently transformed tobacco leaves. Plants expressing ATG8f-mRFP only and leaves transiently co-transformed with free GFP and ATG8f-RFP were used as a negative control. Scale bars: 10 µm (a), 5 µm (b, c).

Considering the previously reported link between ARP2/3 and autophagy, we investigated if the ARP2/3 complex physically interacts with ATG8f. We used plant material co-expressing GFP-tagged ARP2/3 subunits and ATG8f-RFP marker and performed a co-IP experiment. We show here that ARPC2 co-immunoprecipitated with ATG8f in *Arabidopsis* (Figure 5d), and that ARPC5 co-immunoprecipitated with ATG8f in infiltrated tobacco leaves (Figure 5e).

We have stimulated the autophagy and thus the formation of autophagic bodies with NAA treatment (500 nM for 1 h, (Rodriguez et al., 2020) of *arpc2*:GFP-*At*ARPC2 plants and analyzed again the colocalization of the two compartments. The mean number of autophagosomes per confocal section increased significantly after the treatment (Figure 6a). Autophagy stimulation resulted in a decrease in the total number of ARP2/3 dots (Figure 6b), but statistically significantly increased the percentage of ARP2/3 dots colocalizing with autophagosomes (Figure 6c).

**figure 6.**
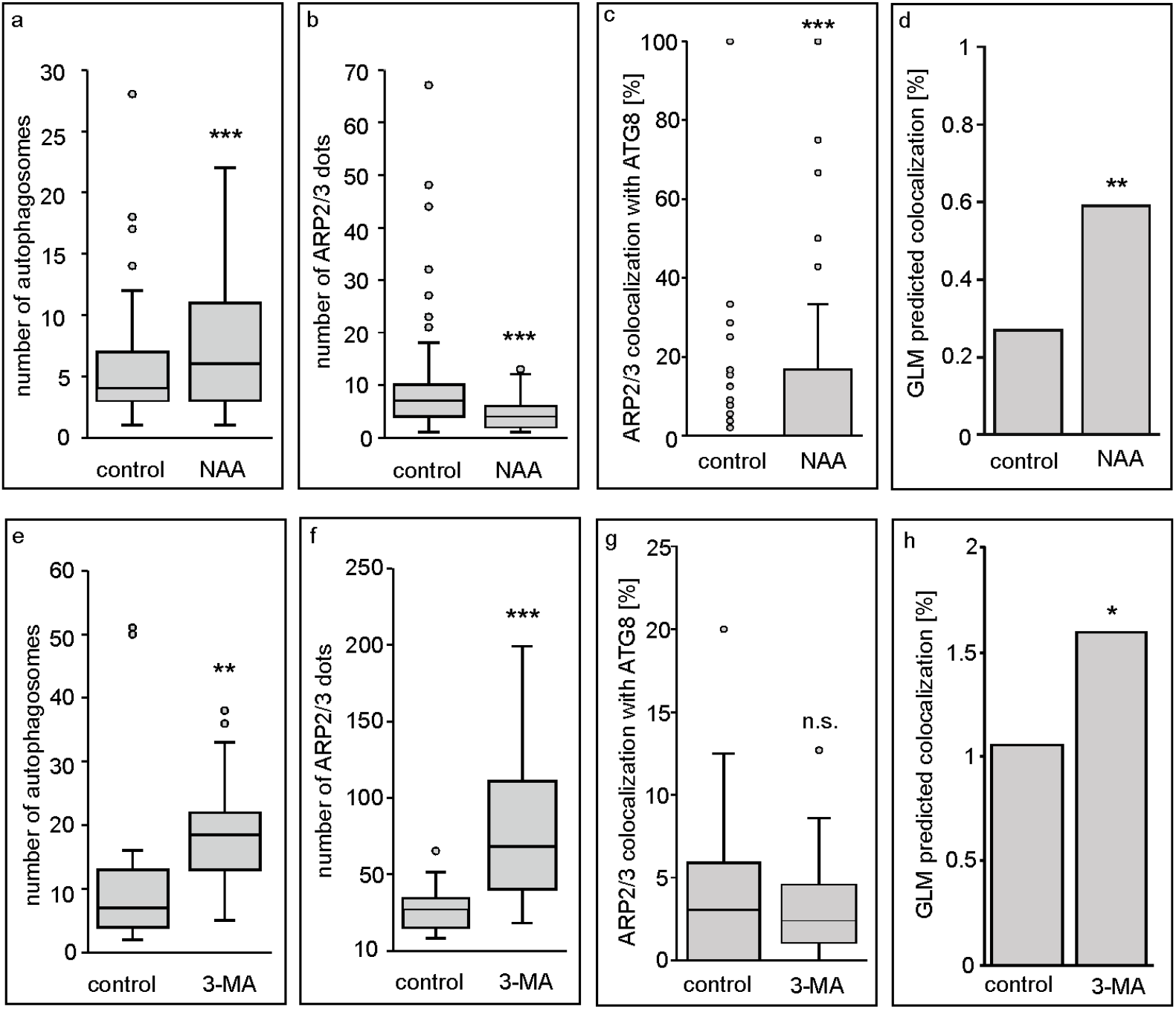
The number of ARP2/3 dots and its colocalization with autophagosomes is sensitive to autophagy induction and inhibition. (**a-c**) 500 nM NAA-induced autophagy in 7DAG *Arabidopsis* seedlings resulted in a significant increase of autophagosomes number detected as ATG8f-mRFP dots (**a**), decreased number of ARP2/3 detected as GFP-*Nt*ARPC2 dots (**b**), and an increased number of the *Nt*ARPC2 dots colocalizing with ATG8f (**c**) in hypocotyl epidermal cells. Particles were calculated manually from confocal microscopy optical sections 106×106 µm through cortical cytoplasm of hypocotyl epidermal cells, using ImageJ. (**d**) The stepwise selection of optimal predictors in the generalized linear model (GLM) proved a significant effect of NAA treatment on increased colocalization. (**e, f**) Autophagy inhibitor 3-MA treatment of 7DAG hypocotyl epidermal cells resulted in a significantly increased number of autophagosomes detected as ATG8f-mRFP dots and an increased number of ARP2/3 detected as GFP-*Nt*ARPC2 dots. (**g**) The number of *Nt*ARPC2 dots colocalizing with ATG8f was not significantly changed when expressed as a percentage of colocalizing *Nt*ARPC2 dots. However, the stepwise selection of optimal predictors in the generalized linear model (GLM) showed a statistically significant effect of 3-MA on increased colocalization of ARP2/3 dots with autophagosomes (**h**). Particles’ numbers were calculated manually from z-stacks of confocal microscopy optical sections 123×123 µm through the cortical cytoplasm of hypocotyl epidermal cells. In **a-d**, for control, n = 127, for NAA treatment, n = 116. In **e-i** for control, n = 19, for 3-MA treatment, n = 22. For **h**, n = 54 (control) and 44 (3-MA). (**d, h**) bar charts represent predicted colocalization after the treatment, based on the GLM with average values for other effects. Statistics represent a chi-square test comparing GLMs without and with the effect of treatment. Statistical significance is indicated by asterisks (*p* < 0.001 = ***, *p* < 0.01 = **, *p* < 0.05 = *, *p* > 0.05 = n.s.)

We have further experimentally interfered with the process of autophagy by applying the autophagy blocker 3-methyladenine (3-MA). 3-MA blocks autophagy in BY-2 tobacco cultured cells (Takatsuka et al., 2004) and 3-MA autophagy block was reported to inhibit starvation-induced peroxisome degradation in BY-2 cells (Voitsekhovskaja et al., 2014). By blocking the autophagy, ARP2/3 dots accumulated in the cytoplasm of both *Arabidopsis* (Figure S6a, b) and BY-2 cells (Figure S6c, d). The effect of the 3-MA autophagy blocker on autophagosomes and ARP2/3 dots accumulation and colocalization was quantified in *Arabidopsis* hypocotyl cells. 10 mM 3-MA treatment resulted in a significant accumulation of autophagosomes (Figure 6e, Figure S6e) and as well as ARP2/3 dots (Figure 6f). Percentage of ARP2/3 dots that colocalized with autophagosomes remained similar compared to the control (Figure 6g). BY-2 cells treated with 5 mM 3-MA also accumulated peroxisomes (Figure S6g). The analysis confirmed that autophagy block by 3-MA induced an increase of peroxisomes (Figure S6g), ARP2/3 dots (Figure S6h) and a proportion of peroxisomes associated with ARP2/3 (Figure S6i). Since the treatment with NAA and 3-MA resulted in multiple effects including the change of particle numbers, we have fitted the data into the generalized linear model (GLM) with the Poisson distribution. The optimal model was selected by stepwise selection according to the Akaike information criterion. The analysis proved that both the NAA and 3-MA treatment statistically significantly increased the colocalization of ARP2/3 with autophagosomes. The significance of the effect of treatment was determined by comparison of full and reduced models using the Chi-square test, *p* < 0.01 (NAA treatment) and *p* < 0.05 (3-MA treatment). In Figures 6d and h, GLM-based prediction of colocalization of ARP2/3 dots with autophagosomes is shown, where only the effect of the treatment is considered. We concluded that autophagy induction and inhibition had an opposite effect on the number of ARP2/3 dots formed in the cytoplasm. Interfering with the autophagy process clearly increased the colocalization of ARP2/3 with autophagosomes and peroxisomes.

### Peroxisomal proteins colocalizing with ARP2/3

The conspicuous localization of proteins in a dot-like peroxisomal peripheral structure has been noted also in other studies. Examples of proteins with such a localization pattern are Snowy cotyledon 3 (SCO3, (Albrecht et al., 2010), Malonyl-CoA decarboxylase (MCD) (Reumann et al., 2009), or effectors of plant fungal pathogen *Colletotrichum higginsianum, Ch*EC51a, and *Ch*EC96 (Robin et al., 2018). To analyze colocalization with ARP2/3 dot, we cloned mCherry-MCD1 and co-expressed it transiently with GFP-*At*ARPC2 in *N. benthamiana* leaves. mCherry-MCD1 protein indeed colocalized with ARP2/3 dots in all observed cells (Figure 7a). Similarly, we confirmed the colocalization of GFP-SCO3 (Figure 7b) and *Ch*EC51a and *Ch*EC96 (Figure 7d,e, respectively) with peroxisomes; all markers colocalized with *At*ARPC5-mCherry on peroxisomes when transiently expressed in tobacco leaves (Figure 7c for SCO3, 7f for *Ch*EC96). These results suggest that the ARP2/3-positive peroxisomal compartment may serve as a general hub for the localization and function of multiple proteins.

**figure 7.**
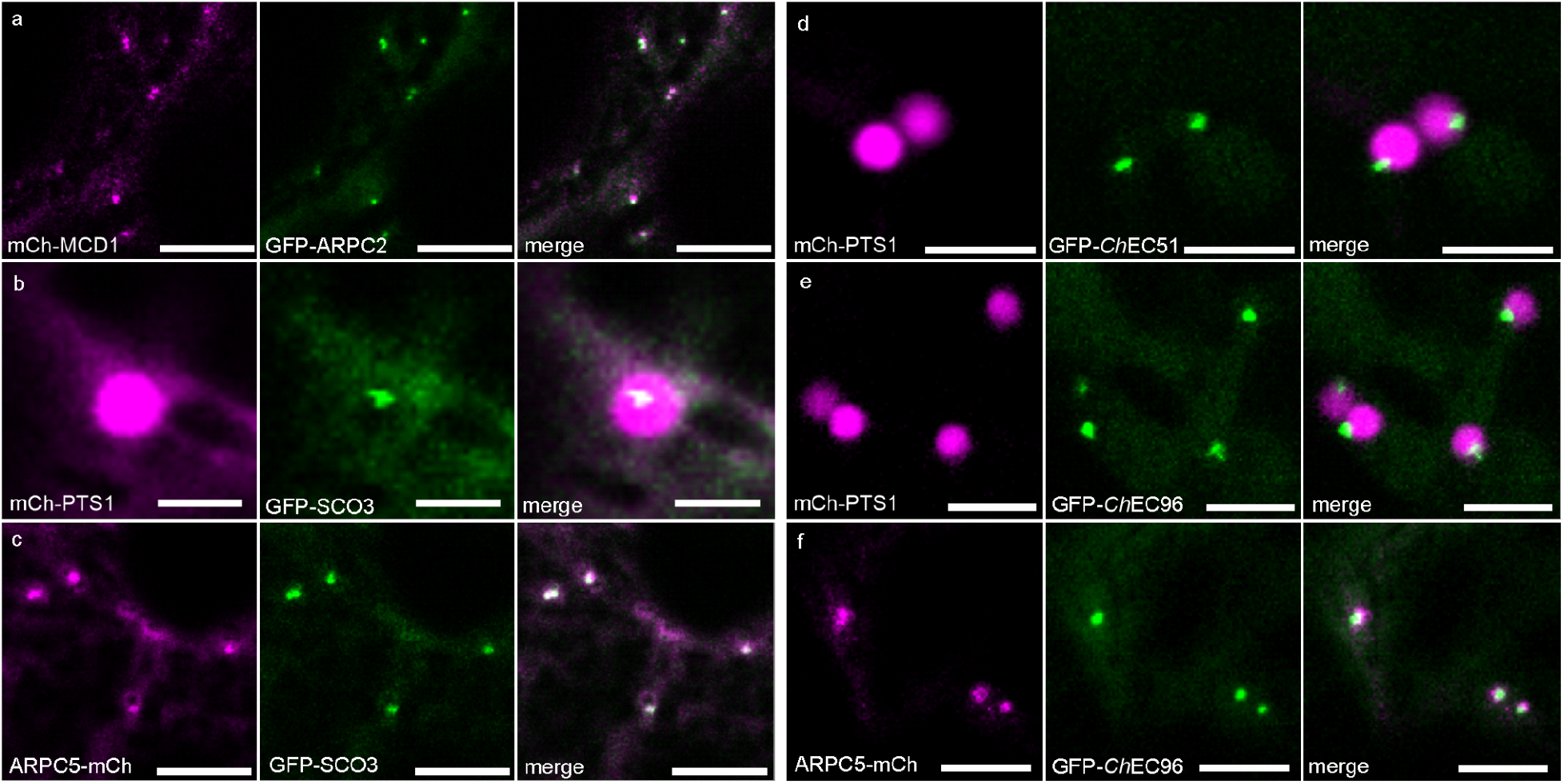
Colocalization of peroxisome-associated ARP2/3 dots with peroxisomal proteins. (**a**) mCherry-MCD1 colocalization with GFP-*At*ARPC2. (b) colocalization of GFP-SCO3 with peroxisomes as a dot-like structure, (c) colocalization of ARPC5-mCherry with GFP-SCO3, (d) colocalization of GFP-*Ch*EC51 and (e) colocalization of GFP-*Ch*EC96 with peroxisomes as dot-like structures, (f) colocalization of *At*ARPC5-mCherry with GFP-*Ch*EC96. All markers colocalized with ARP2/3 on peroxisomes. Scale bars = 10 μm (a, c) 2 μm (b), 5 μm (d-f), transient expression in leaf epidermis of *N. benthamiana*.

## DISCUSSION

In this work, we studied the localization of ARP2/3 in plant cells expressing complex subunits ARPC5 and ARPC2 fused to fluorescent proteins. *Arabidopsis* subunits GFP-*At*ARPC2 and GFP-*At*ARPC5, and tobacco subunit GFP*-Nt*ARPC2 formed a motile dot-like structure in the cytoplasm, which is consistent with our previous observations (Havelková et al., 2015). Both FP-fused subunits showed also a diffuse cytoplasmic signal. A motile dot-like localization of both subunits was detectable in all tested cell types, and dot-like structures were formed also in cells with very low expression (β-estradiol-inducible GFP-*Nt*ARPC2 induction with 2 nM β-estradiol), and our vectors rescued distorted trichomes phenotypes. This suggests that ARP2/3 dots are native structures.

Based on the following evidence, we believe that the ARP2/3 dot comprises the whole and activated complex. First, when we expressed *At*ARPC5-mCherry and GFP-*At*ARPC2, both subunits localized to the same structure. Second, NAP1, a subunit of the ARP2/3-activating complex (Zimmermann et al., 2004), colocalized with peroxisomes in an identical way as ARP2/3 subunits, confirming that also activation complex is part of the ARP2/3-positive peroxisome-associated structure. Third, the three analyzed components of ARP2/3 and WAVE/SCAR activation complex colocalized and co-immunoprecipitated with each other in the GFP pull-down assay. Fourth, the GFP-pull down experiment followed by protein identification using MS/MS allowed us to identify all ARP2/3 subunits as interactors. In contrast, WAVE/SCAR regulatory complex subunits were not identified among interacting proteins. This could be explained by the transient nature of ARP2/3 and WAVE/SCAR interaction.

If the ARP2/3 peroxisome-associated dot contains an assembled and active ARP2/3 complex, we expect the ARP2/3 complex to perform its conserved role there, which is the nucleation of actin filaments. Surprisingly, we found limited evidence that actin filaments are related to this structure. In living cells, ARP2/3 dot turning around the motile peroxisome had no clear and stable interaction with fluorescently labeled actin cables, along which the organelle is transported (Mano et al., 2002; Mathur et al., 2002). This is consistent with the fact that peroxisomes’ motility was not changed in several ARP2/3 mutant lines. Therefore, the ARP2/3 structure on peroxisome probably does not play a role in organelle movement along actin. However, we show here that ARP2/3 dot colocalized with actin, which was detectable in fixed cells stained by phalloidin, a small molecule with a high affinity to polymerized actin (Blancaflor, 2000). Indeed, we observed diffuse actin stained by phalloidin-TRITC surrounding the GFP-*Nt*ARPC2 signal in fixed cells. We confirmed these results in co-IP experiments using GFP-tagged ARP2/3 subunits that pulled down actin in total protein extracts. This raises a possibility that ARP2/3 dots associated with peroxisomes interact with, or nucleate a very fine actin network there. Thin and branched actin filaments, which are of a highly dynamic nature, may not be detectable using plant actin *in vivo* markers, which are fragments of associated proteins (FABD, (Ketelaar et al., 2004), or peptides binding polymerized actin (Lifeact, (Riedl et al., 2008) fused to fluorescent proteins. Actin visualization in ARP2/3 peroxisomal dots clearly requires methods providing images with a resolution higher than fluorescence microscopy, which is, for example, electron microscopy.

Our experiments suggest that intact actin cytoskeleton is not required for ARP2/3 association with peroxisome, because ARP2/3 localization was not perturbed upon latrunculin B treatment. However, it seems that the formation of the dot-like structure requires the presence of one of the core subunits, ARPC2 (Robinson et al., 2001; Kotchoni et al., 2009), while the presence of the peripherally located ARPC5 subunit is not needed for the structure formation. Interestingly, the complex break-up in *arpc2* mutants resulted in the loss of peripheral localization and GFP-ARPC5. The analysis using Airyscan superresolution microscopy confirmed the localization of ARP2/3 at the periphery of the organelle. ARP2/3 is a cytoplasmic complex, which is in concordance with our observation of a diffuse cytoplasmic signal in cells expressing ARP2/3 markers fused to GFP or RFP. It is expected that the ARP2/3 dot is localized on the cytoplasmic side of the peroxisome, where it performs its actin-related functions. Indeed, our inspection of sequences confirmed that none of the ARP2/3 subunits contains any known peroxisome-localization sequence (data not shown). Dispersed localization of GFP-ARPC5 in *arpc2* mutants may be caused by the loss of the ability to assemble the full complex.

To understand the function of ARP2/3 in peroxisomes, we analyzed the peroxisomal structure and functions in mutants lacking ARP2/3 complex. We show here that basic metabolic functions of ARP2/3 mutant peroxisomes such as β-oxidation of fatty acids or IBA conversion to auxin are not affected. Our results demonstrating a higher number of peroxisomes in mutant lines point to defects in the control of peroxisome division, biogenesis or degradation. Although the division of peroxisomes was often observed in our experiments, ARP2/3 dots on peroxisomes did not localize to the division sites, which excludes a possible role of ARP2/3 in the division. We focused on the possible role of ARP2/3 in peroxisome degradation, called pexophagy (Farmer et al., 2013; Lee et al., 2014), and we show here a strong link between peroxisome-localized ARP2/3 and autophagy pathway. Actin polymerization participates in autophagy processes in yeast and mammalian cells (reviewed in (Coutts and La Thangue, 2016; Kast and Dominguez, 2017). ARP2/3 complex has been demonstrated to be involved in autophagy in several recent studies, including the relationship of this specific ARP2/3 function to pathogenesis in some diseases (Xia et al., 2013; Zavodszky et al., 2014; Coutts and La Thangue, 2015; Kast et al., 2015; Kast and Dominguez, 2015; Mi et al., 2015; Mathiowetz et al., 2017; Rivers et al., 2020; Sarkar et al., 2021). In plants, ARP2/3 has been shown to participate in processes that link autophagy and endocytosis (Wang et al., 2016, 2019). A specific type of autophagy securing the degradation of peroxisomes engages ARP2/3-dependent polymerization in yeast (Monastyrska et al., 2008). In our experiments, we found a low frequency of ARP2/3 peroxisomal dot and autophagosome marker ATG8f colocalization. Microscopic analysis revealed that the ARP2/3 and ATG8f association was often of transient nature, where the two structures were attaching and detaching during the time-lapse acquisition. This observation is consistent with brief colocalization events between Atg8 and Arp2 in yeast (Monastyrska et al., 2008). Thus, a rather low frequency of colocalization may be explained by the dynamic and transient nature of the ARP2/3 peroxisomal dot and ATG8f marker interaction. Despite limited colocalization of ARP2/3 peroxisomal structure and autophagosome, our results indicate a strong relationship between ARP2/3 and pexophagy. First, we observed that if both markers were present on the peroxisome, they always colocalized. Second, the co-IP experiment proved ATG8f-RFP and

ARP2/3 subunits physical interaction. Third, interfering with autophagy processes resulted in changes in the number of ARP2/3 dot structures. Activation of autophagy by NAA resulted in a significant decrease in the number of ARP2/3 dots, while blocking of autophagy by 3-MA significantly increased their amount. ARP2/3 role in autophagosome formation or function in plant peroxisomes is not clear at the current stage of the research; only after specifying its role, we can explain increased rates of autophagosome and ARP2/3 colocalization after both NAA and 3-MA treatment. More details about the pexophagy process are also needed to explain our observation that the ATG8f marker was predominantly observed as a small dot associated with peroxisome, while advanced steps of pexophagy with peroxisome engulfed with autophagosome membrane were observed very rarely. Blocking of autophagy with 3-MA that induced a dramatic increase of peroxisomal numbers and rate of peroxisomes with ARP2/3 dot in BY-2 cells suggests that the increase in peroxisomal numbers that were detected in *Arabidopsis* ARP2/3 mutants were caused by a delay in peroxisome degradation upon non-functional ARP2/3 complex.

The presumable role of actin in autophagy is the facilitation of membrane remodeling by ARP2/3-generated branched actin reviewed in (Coutts and La Thangue, 2016; Kast and Dominguez, 2017). Localization of ARP2/3 with ATG8f marker at the surface of peroxisomes is consistent with this. Fine and dynamic polymerized actin generated by ARP2/3 may also play a general role in plant peroxisomes. Note that many peroxisomes contained only ARP2/3 dots without the ATG8f marker. Further, a number of peroxisome enzymes were identified as interactors of ARP2/3 in our experiment, although we did not observe problems with β-oxidation in mutant plants. As shown by super-resolution STED microscopy in fibroblasts, peroxisomal membrane proteins are not homogeneously distributed on the peroxisome periphery, but they are organized into subdomains (Galiani et al., 2016). In plant peroxisomes, several peroxisomal proteins and enzymes are not distributed homogeneously in the lumen of peroxisomes but localize to a discrete peroxisomal sub-compartment overlapping only partially with the luminal marker of peroxisomes. Examples of such proteins are Malonyl-CoA decarboxylase and Indigoidine synthase A (Reumann et al., 2009) or unknown protein 9, supposedly a peroxisomal lipase (Quan et al., 2013). Non-homogenous peroxisomal localization of peroxisome proteins Hydroxy-acid oxidase 1, Harmless to ozone layer 3, or trimmed TSK-associating protein 1 was reported by Pan (Pan et al., 2018). Interestingly, newly identified weak peroxisome localization peptide SKV, when fused to YFP, was also localized to dot structures in peroxisomes (Reumann et al., 2009). A recent study suggests that peroxisomes may form intraluminal vesicles, which can also contribute to the non-homogeneous distribution of luminal and membrane peroxisomal proteins (Wright and Bartel, 2020). In our work, we tested the localization of proteins SNOWY COTYLEDON3 (Albrecht et al., 2010), MCD (Reumann et al., 2009), and *Colletotrichum higginsianum* effectors *Ch*EC51a and *Ch*EC96 (Robin et al., 2018) with ARP2/3. In all cases, we observed a perfect colocalization with ARP2/3 peripheral peroxisomal structure. The localization of several peroxisome-targeted enzymes, identified as possible ARP2/3 interactors in our proteomic study, has not been tested yet.

Since membrane remodeling by polymerization of branched actin filaments is a conserved role of ARP2/3, we suggest that the ARP2/3-generated actin filaments form a scaffold facilitating membrane-related processes in the peroxisome periphery. These could include protein compartmentation within the peroxisomal membrane including substrates for ATG8f binding and degradation during peroxisome degradation.

## METHODS

### Plant material and cultivation

*Arabidopsis thaliana* genotypes used in this study were Col-0 (wild-type), *arpc5* (SALK_123936.41.55) and *arpc2/distorted2-1* (El-Din El-Assal et al., 2004). For IBA treatment, T-DNA insertion mutants *arp2* (SALK-077920) and *arpc4* (SALK-013909) have been employed (Sahi et al., 2018). For *in vitro* cultivation, seeds were briefly washed with 96% ethanol and then surface-sterilized by bleach solution (sodium hypochlorite solution at the final concentration of 2.5 %) with 0.01% Triton X-100 for 10 minutes, washed in sterile H_2_O and sown on vertical plates with agar medium (1 % [w/v] sucrose, 2.2 g/l MS salts [Sigma/Aldrich], 0.8 % agarose, [pH 5.7]) in cultivation room under a photoperiod of 16 h light:8 h darkness and 23 °C and light intensity 110 µmol/m^2^/s.

Cells of tobacco line BY-2 (*Nicotiana tabacum* L. cv Bright-Yellow 2) (Nagata et al., 1992) were cultivated in darkness at 26 °C on an orbital incubator (120 rpm, orbital diameter 30 mm) in a liquid medium (3% [w/v] sucrose, 4.3 g/l Murashige and Skoog salts, 100 mg/l inositol, 1 mg/l thiamin, 0.2 mg/l 2,4-dichlorophenoxyacetic acid, and 200 mg/l KH_2_PO_4_ [pH 5.8]) and subcultured weekly. Stock BY-2 calli were maintained on media solidified with 0.6% (w/v) agar and subcultured monthly. Transgenic cells and calli were maintained on the same media supplemented with 100 μg/ml kanamycin or 20 μg/ml hygromycin. All chemicals were obtained from Sigma-Aldrich unless stated otherwise.

*N. benthamiana* plants for transient leaf transformation using the *Agrobacterium* infiltration technique were grown in peat pellets in a growth chamber under a photoperiod of 16 h light:8 h darkness and 23 °C and light intensity 110 µmol/m^2^/s.

For expression of constructs with β-estradiol-inducible system XVE, the medium was supplemented with 2nM - 10μM β-estradiol.

### In vivo markers

Subunits *At*ARPC2, *At*ARPC4 and *At*ARPC5 were amplified from *Arabidopsis* cDNA using primers with restriction sites overhangs (Supporting Table S2). Restriction enzymes *BamHI* and *HindIII* were used for cloning the fragment to the pGreen vector adapted for N-terminal GFP fusion, with the 35S promoter. For the production of *At*ARPC5-mCherry, restriction sites for *BamHI* and *XmaI* were used, and the fragment was cloned to the *p*Green vector as a C-terminal fusion with the mCherry fluorescent protein under the 35S promoter. Further, a vector with a β-estradiol-inducible XVE system (Zuo et al., 2000) was used for the expression of GFP-*Nt*ARPC2 (Havelková et al., 2015). MCD (Malonyl-CoA decarboxylase) sequence was amplified with PCR from *Arabidopsis* cDNA using primers with restriction sites overhangs (Supporting Table 2). *BamHI* and *EcoRI* enzymes were used to clone the fragment to the pGreen vector as a C-terminal fusion with mCherry, under the 35S promoter.

Following marker lines were used as well: mCherry-PTS1, GFP-PTS1 and CFP-PTS1 for peroxisomes visualization, (Nelson et al., 2007), ATG8f-mRFP for autophagosome visualization (Honig et al., 2012), GFP-FABD (Voigt et al., 2005) and Lifeact-mRFP (Fendrych et al., 2013) for actin visualization, GFP-SCO3 (Albrecht et al., 2010), *Colletotrichum higginsianum* effectors *Ch*EC51a and *Ch*EC96 fused with GFP (Robin et al., 2018), and NAP1-GFP expressed under native promoter in *nap1* mutant (Wang et al., 2016).

### Transformation of plant material

*Arabidopsis* plants were stably transformed by *Agrobacterium tumefaciens* strain C58C1 or GV3101 using the floral dip method (Zhang et al., 2006). Transformed seedlings were selected on agar plates with antibiotics. FAST transformation technique based on the co-cultivation of *Arabidopsis* seedlings with *A. tumefaciens* suspension as described in (Li et al., 2009) was used for transient transformation of young *Arabidopsis* seedlings with the following modifications. First, 25 µl/l of Silwet L-77 was used during seedlings co-cultivation with *Agrobacterium*. Second, 100 mg/l cefotaxime instead of sodium hypochlorite was added to the washing solution for the wash-out of the *Agrobacterium*. BY-2 cells were transformed by co-cultivation with *Agrobacterium tumefaciens* as described in (Klíma et al., 2019). *N. benthamiana* leaves were transiently transformed using the *Agrobacterium* infiltration technique. briefly, the overnight culture of transgenic *Agrobacterium* was centrifuged (5 min, 3500g) and washed three times with infiltration solution (10 mM MgCl_2_, 10 mM MES, pH 6.5), resuspended in infiltration solution to obtain OD_600_ = 0.5 and supplemented with 200 μM 4′-Hydroxy-3′,5′-dimethoxyacetophenone (acetosyringone) and cultivated for 3 hours. The bacterial suspension was then injected into the mesophyll through the abaxial leaf epidermis using a 5ml syringe. Infiltrated tobacco plants were cultivated for 2 days and then observed using a confocal microscope.

### Fixation of plant material

For TRITC-phalloidin staining, cells were fixed in actin stabilizing buffer (ASB – 100 mM PIPES, 4 mM MgCl_2_, 10 mM EGTA, pH = 6.9) supplemented with 400 μM MBS (N-succinimidyl 3-maleimide benzoate) for 20 minutes and stained in ASB supplemented with 0.3 M mannitol, 2% glycerol and 0.1 µM TRITC-phalloidin (Sigma-Aldrich, P1951) for 5 min. After washing in the ASB buffer, cells were immediately observed. For fixation of plant material expressing fluorescent markers, plantlets or cells were fixed in ASB supplemented with 400 μM MBS and 3.2% formaldehyde for 30 minutes. Plants or cells were observed after washing in PBS.

### Clearing of plant material

For the microscopic examination of mutant trichomes we incubated true leaves of *Arabidopsis* in a clearing solution (120 g chloral hydrate, 50 ml water, 7,5 ml glycerol) at room temperature until cleared (depending on the size of the sample, approximately for 2 days).

### Microscopy

Confocal laser scanning microscopy was performed using Zeiss TCS 880 and Leica SP8. For Zeiss TCS 880, confocal pictures were taken using C-Apochromat 40x magnifying water immersion objective, NA = 1.2, and predefined excitation and emission wavelength for CFP, GFP and mCherry, as provided by Carl Zeiss ZEN Black microscope operating software (excitation 488 nm and emission 495-530 nm for GFP, excitation 561 nm and emission 581-632 nm for mCherry and excitation 405 nm and emission 415-475 nm for CFP). For the simultaneous acquisition of CFP, GFP and mCherry (Figure 5b, c), the Linear unmixing feature was used in Zeiss ZEN Black software. For Leica SP8, water immersion objective HC PL APO CS2 63x with NA = 1.2. Excitation 488 nm and emission 490-550 nm (GFP), excitation 561 nm and emission 580-650 nm (mCherry). The pinhole was set to 1AU at 580 nm.

Confocal spinning disc microscopy was performed using Nikon Ti. Images were taken using Plan Apo 100x oil immersion objective, NA = 1.45 and camera Andor Zyla VSC-01592. Excitation laser 488 nm with emission filter 525/30, and excitation laser 561 nm with emission filter 600/52 were used for GFP and mCherry fluorescent proteins, respectively.

For Airyscan superresolution microscopy, 5-day-old seedlings of plants expressing GFP-*Nt*ARPC2 were fixed for 20 min in ASB supplemented with 400 μM MBS. Inverted confocal laser scanning microscope Zeiss LSM880 (Carl Zeiss, Jena, Germany) equipped with alpha Plan-Apochromat 100×/1.46 Oil objective and Airyscan detector was used. GFP (ex: 488nm, em: 495-550) and RFP fluorescence channels (ex: 561nm, em: 555-620) were acquired with the SR mode of Airyscan detector in a sequential scanning setup. Airyscan Processing of raw files was performed in the ZEN black software (Carl Zeiss, Jena, Germany). Microscopy data were analyzed in the ZEN blue 2.5 software (Carl Zeiss, Jena, Germany) where 3D Transparency rendering and 3D surface reconstruction mode were used.

### Quantification of peroxisomal phenotypes

For peroxisomes visualization and quantification, plants expressing GFP-PTS1 and mCherry PTS1 were used. Confocal optical sections 60 × 60 µm from the center of an adaxial side of the cotyledon epidermis were obtained. The number of peroxisomes per confocal section was measured using a Multi-point tool in ImageJ software. The numbers of peroxisomes were compared between mutants and WT by a T-test. To measure the size of peroxisomes, individual peroxisomes from confocal sections were selected using the Wand tool in ImageJ software and the area of individual peroxisomes was measured, mutants and WT were compared using a T-test. The average velocity of peroxisomes in WT, *arpc2* and *arpc5* mutants were measured in the cortical region of hypocotyl epidermal cells in *Arabidopsis thaliana* seedlings from confocal microscopy time series (60 frames, Δt = 1.29 s) using TrackMate plugin in ImageJ software (Tinevez et al., 2017) and compared between WT and mutants using a T-test. For each measurement, two or three biological replications were done, with tens of plants per replicate.

The data with skewed distribution were normalized using a square root transformation before T-test.

### Quantification of ARP2/3 dot-autophagosome association

To quantify the association of ARP2/3 dot with autophagosomes, we used *arpc2* plants expressing GFP-*Nt*ARPC2 and autophagosome marker ATG8f-mRFP. Total numbers of autophagosomes and ARP2/3 dots, as well as the number of ARP2/3 dots in association with autophagosomes, was counted in fixed plant material. The percentage of colocalizing particles was expressed as a percentage of ARP2/3 dots colocalizing with autophagosomes from a total number of ARP2/3 dots. The differences between control and treatment were compared using a T-test. The data with skewed distribution were normalized using a square root transformation before T-test. For evaluation of complex effects of NAA and 3-MA treatment, the generalized linear model (GLM) with Poisson distribution was fitted to the data in the R (R Core Team, 2021) and the optimal combination of predictors was determined by stepwise selection based on AIC. Statistical significance of the treatment effect was determined by chi-square test.

### Isolation of proteins and co-immunoprecipitation

For total protein fraction isolation, approximately 1g of infiltrated tobacco leaves or 10-day-old *Arabidopsis* seedlings were frozen in liquid nitrogen and homogenized with mortar and pestle. The frozen powder was mixed 1:1 (v/w) with 2x concentrated MES buffer (25 mM MES; 5 mM EGTA; 5 mM MgCl_2_; 1 M glycerol; pH = 6.9) supplemented with a protease inhibitors cocktail (Sigma-Aldrich, P9599) and was let thaw on ice. The sample was then centrifuged at 3000 g for 15 min at 4 °C and the supernatant was centrifuged at 25000 g for 30 min at 4 °C. The supernatant contained 2.5 mg proteins/ 1 ml (determined using Bio-Rad Protein Assay, Cat. No. 5000006). This protein fraction was used for co-immunoprecipitation (input). 1 ml of protein extract was mixed with 50 µl of magnetic beads from µMACS™ GFP Isolation Kit (Miltenyi Biotec) and the mixture was incubated on ice while mild shaking. The extract was then loaded into the column in a magnetic stand and GFP-tagged proteins and their interactors were isolated according to the manufacturer protocol. Isolated immunoprecipitated proteins (the whole amount of eluted fraction, 50 µl) as well as respective inputs (20 µl of extracts corresponding to 50 µg of total proteins) were separated using SDS PAGE electrophoresis and transferred onto nitrocellulose membrane by electro-blotting (SemiDry, BioRad). Western blots were probed with rabbit anti-GFP (1:5000; Agrisera AS152987), rabbit anti-RFP (1:6000; Abcam, ab167453) and anti-actin (1:4000; MP, 0869100-CF).

For peroxisomal protein fraction isolation, protoplasts were prepared from 4-day-old BY-2 cells expressing mCherry-PTS1 and estrogen-inducible GFP-*Nt*ARPC2 by the cultivation of cells in a protoplasting medium (400 mM mannitol, 20 mM MES, 1 mM EDTA, pH = 7.5, supplemented with 1% cellulase and 0.1% pectolyase Y-23) for 3 hours by slow shaking (50 rpm). Protoplasts were shredded with Dounce homogenizer on ice. The homogenized sample was supplemented with a protease inhibitor cocktail and layered on the top of the pre-cooled sucrose gradient, consisting of 15%, 40% and 60% sucrose. The gradient was centrifuged at 31000 g for 1 hour at 4 °C. The peroxisome-rich fraction was found on the 40%-60% sucrose interface, and the presence of peroxisomes and ARP2/3 dots was confirmed by fluorescence microscopy. Free GFP and GFP-tagged proteins were subsequently extracted from the peroxisome-rich fraction using the µMACS GFP Isolation Kit as described above. Isolated proteins were analyzed by nLC-MS 2 analysis (see Supporting data for details).

## ACKNOWLEDGEMENTS

Acknowledgement to Dr. Karel Harant and Dr. Pavel Talacko (Laboratory of Mass Spectrometry, Charles University, Faculty of Science) for proteomic and mass spectrometric analysis. The work was supported by the Ministry of Education, Youth and Sports of the Czech Republic, project No. 19-10845S and Grant Agency of the Charles University No. 816217. It was supported by Leverhulme Trust funding (RPG-2015-106). Microscopy was performed in the Laboratory of Confocal and Fluorescence Microscopy co-financed by the European Regional Development Fund and the state budget of the Czech Republic, projects no. CZ.1.05/4.1.00/16.0347 and CZ.2.16/3.1.00/21515 and supported by the Czech-BioImaging large RI project LM2018129. Computational resources were supplied by the project “e-Infrastruktura CZ” (e-INFRA LM2018140) provided within the program Projects of Large Research, Development and Innovations Infrastructures. Superresolution microscopy was performed in the Imaging Facility of the Institute of Experimental Botany CAS, which was supported by the MEYS CR (Large RI Project LM2018129 Czech-BioImaging). We would like to thank Matyáš Fendrych for many useful comments and suggestions to the manuscript.

## SHORT LEGENDS FOR SUPPORTING INFORMATION

### Supporting Data

Supporting Data 1 describes protein digestion and nLC-MS^2^ analysis details.

### Supporting Tables

Supporting Table 1 contains a list of consensus protein interactors of GFP-*Nt*ARPC2 in peroxisome-enriched protein fraction, identified in both repetitions of proteomic analysis.

Supporting Table 2 contains a list of primers used in this study.

### Supporting movie

https://drive.google.com/drive/folders/1GZrpslcMmi2i8ZCJjhmPHQq3VD6gSxP9?usp=sharing

Movie S1 shows motility of peroxisomes (mCherry-PTS1, magenta) with ARP2/3 dot (GFP-*Nt*ARPC2, green) in hypocotyl cells of transgenic *Arabidopsis thaliana*.

Movie S2 shows motility of peroxisomes (mCherry-PTS1, magenta) with ARP2/3 dot (GFP-*Nt*ARPC2, green) along actin filaments (FABD-GFP, green) in pavement cells of transgenic *Arabidopsis thaliana*.

Movie S3 shows transient and occasional colocalization of peroxisome-associated ARP2/3 dot (GFP-*Nt*ARPC2, green) with autophagosome marker ATG8f (magenta).

## SUPPLEMENTARY FIGURES

**figure S1.**
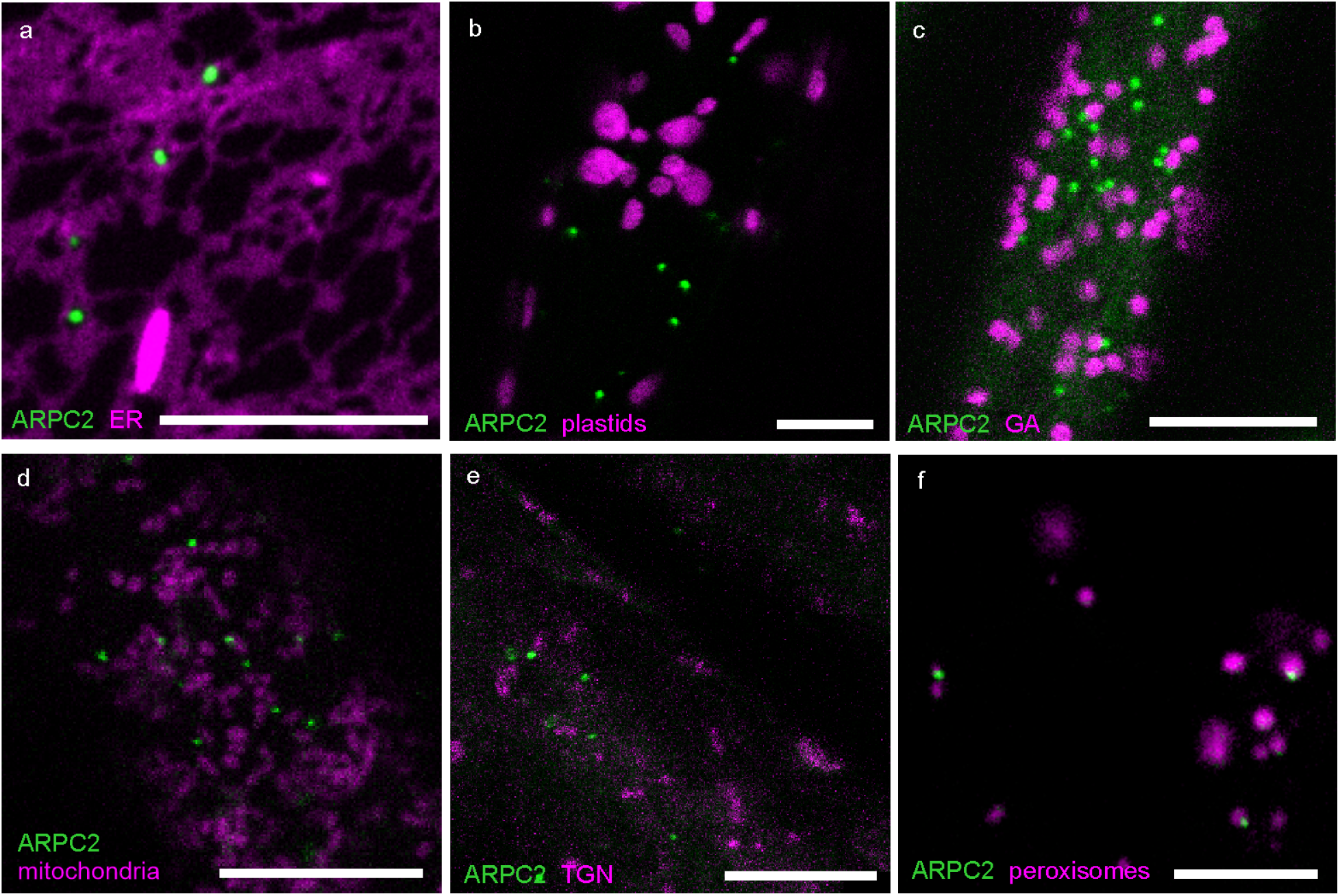
Co-expression of GFP-*Nt*ARPC2 with different organellar markers shows the exclusive association of ARP2/3 dots with the peroxisomes. Colocalization was observed in *Arabidopsis thaliana* hypocotyl epidermal cells (a, b, c, e, f) and in BY-2 tobacco suspension cells (d) expressing GFP-*Nt*ARPC2 with co-visualized organelles. (a-e) The ARP2/3 dots do not colocalize with ER, visualized by co-expressed ER-RFP marker (a, Nelson et al., 2007), plastids (b, chlorophyll autofluorescence), co-expressed GA-RFP marker (c, Nelson et al., 2007), mitochondria labelled by MitoTracker Red (d,) or TGN, visualized by co-expressed RABD2A-RFP marker (e, Geldner et al., 2009). (f) ARP2/3 dots are exclusively associated with a subpopulation of peroxisomes, visualized by mCherry-PTS1 marker Nelson et al., 2007). Scale bars = 10 µm.

---

**figure S2.**
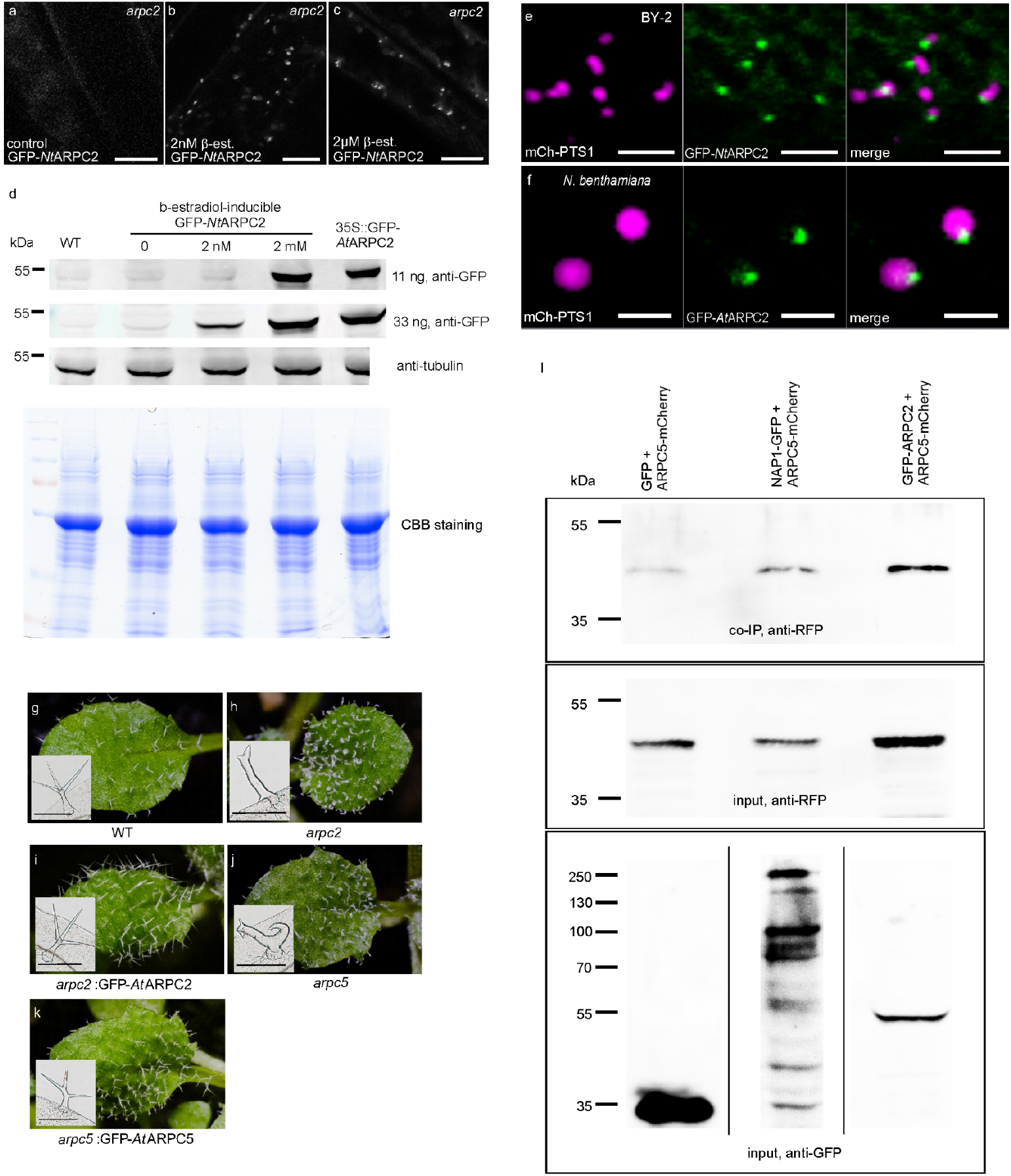
(a-c) Motile dots formed by inducible GFP-*Nt*ARPC2 were observed under conditions of very low protein expression (b, 2 nM β-estradiol, compare with c, 2 µM β-estradiol), while no signal was observed in uninduced controls (a). (d) Detection of GFP-*Nt*ARPC2 in Western blots demonstrates the expression levels of the GFP-*Nt*ARPC2 subunit induced by 2 nM and 2 µM β-estradiol in cells shown in (a-c). Note that 2 nM β-estradiol-induced cells have considerably lower expression, which is undetectable if 11 ng of total proteins is loaded when compared to cells expressing under the 35S promoter or cells induced by 2 µM β-estradiol. Beta-tubulin levels are shown as a control of protein loading. (e, f) GFP-*Nt*ARPC2 and GFP-*At*ARPC2 motile dots colocalize with peroxisomes in stably expressing BY-2 cells (e) and transiently transformed tobacco leaf pavement cells (f). (g-k) The expression of GFP-fusion proteins rescued the phenotype of distorted trichomes of *arpc2/dis2-1* (h) and *arpc5* (j) mutant line. True leaves of mutants expressing respective fusion protein (i, k) resembled WT trichomes (g). (i) ARP2/3 subunits GFP-ARPC2 and ARPC5-mCherry, and ARP2/3 subunit ARPC5-mCherry and NAP1-GFP co-immunoprecipitate in extracts from *Arabidopsis* stably expressing respective markers. Scale bars: (a-c) = 10 µm, (e, f) = 2 µm, (g-k) = 200 µm.

**figure S3.**
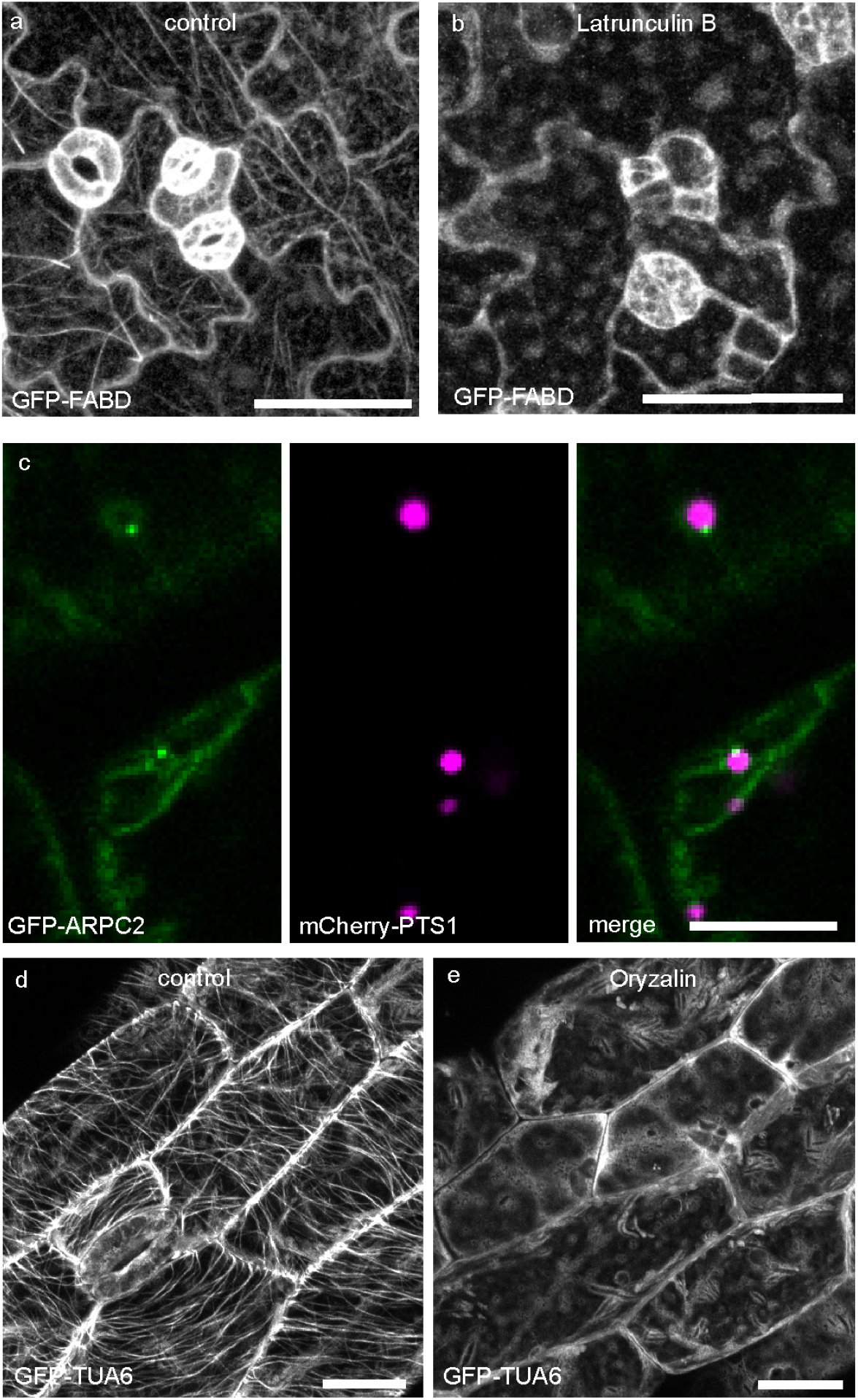
(a, b) Actin filaments marked by GFP-FABD in *Arabidopsis* cotyledons were completely depolymerized after 30 min of treatment with 500 nM latrunculin B. (c-e) Treatment with 40 mM oryzalin for 3 hours did not alter the structure or peroxisomal localization of ARP2/3 dots (c), although it dissolved the microtubular cytoskeleton completely (f, compared to untreated control in d). Scale bars: 50 µm (a, b), 20 µm (c, d, e).

**figure S4.**
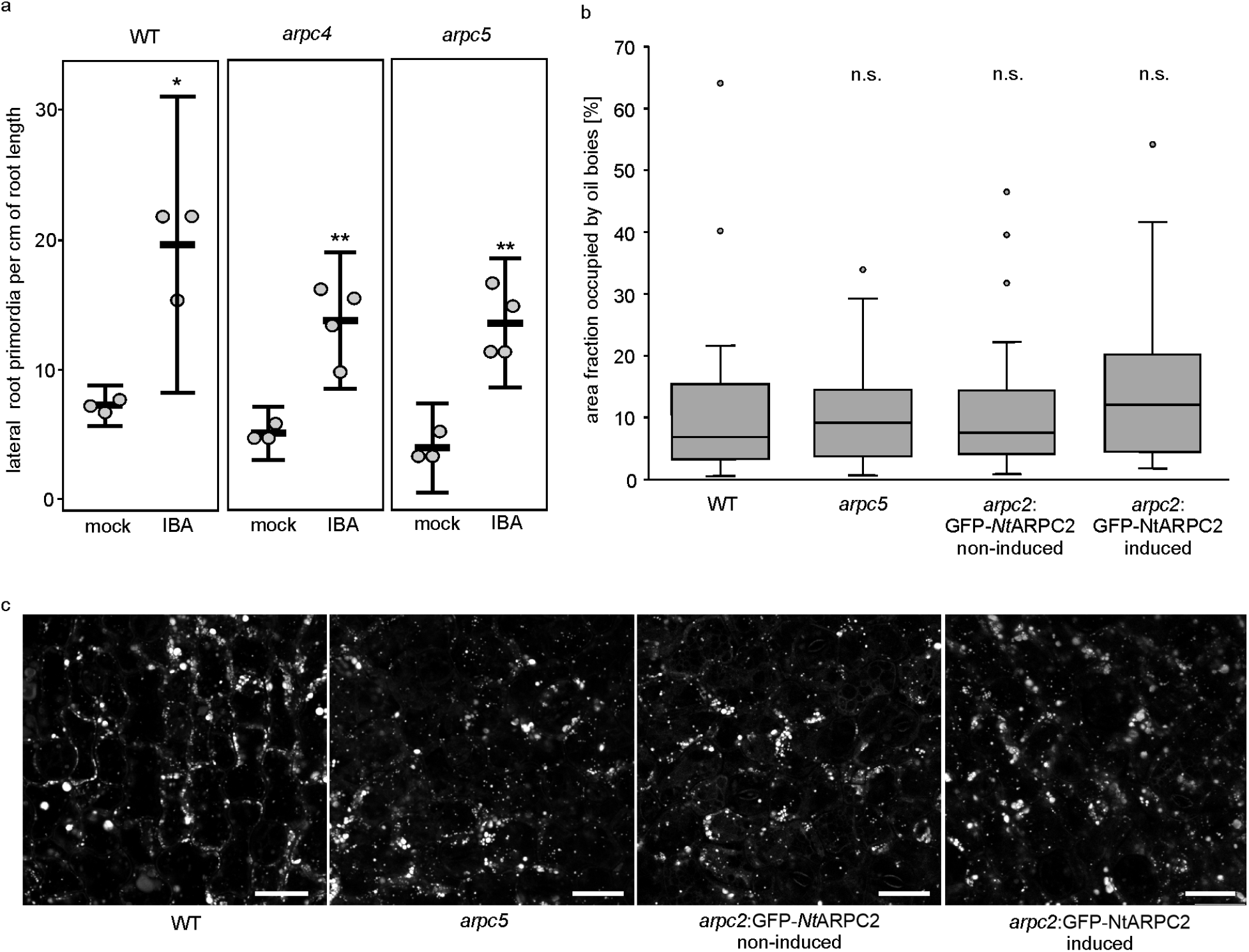
(a) ARP2/3 mutant lines *arpc4* and *arpc5* responded to IBA by increased initiation of lateral roots, indicating a functional β-oxidation pathway that converts IBA to auxin. Central lines indicate the mean, error bars indicate 95% CI. Significance of the T-test are indicated by asterisks (*p* < 0.05 = *, *p* < 0.01 = **). (b, c) The degradation of storage oil bodies in ARP2/3 mutants was not significantly different when compared to WT. Area fraction occupied by oil bodies was calculated for confocal sections, n = 20 (WT), 19 (*arpc5*), 20 (non induced *arpc2*:GFP-*Nt*ARTPC2), 20 (induced *arpc2*:GFP-*Nt*ARPC2); 10 plant were analyzed in each variant. Central lines in boxplots represent the median. T-tests comparing WT and each mutant showed no significant difference (n.s., *p* > 0.05). Scale bars in c = 20 µm. Oil bodies quantification: For quantification of oil bodies degradation, 4-days-old dark-grown seedlings on MS medium without sucrose were stained with Nile red. The stock solution of Nile Red (0.5 mg/ml in acetone) was diluted at 1:1000 in distilled water. The seedlings were incubated in this solution at room temperature for 30 minutes. Oil bodies were imaged in the etiolated hypocotyl epidermal cells using a confocal microscope (561 nm laser excitation, 575-630 nm emission). For quantification of oil bodies amount in the cells, the image area occupied by oil bodies in the single optical frame, that comprised several cells, was measured using the threshold function in ImageJ.

**figure S5.**
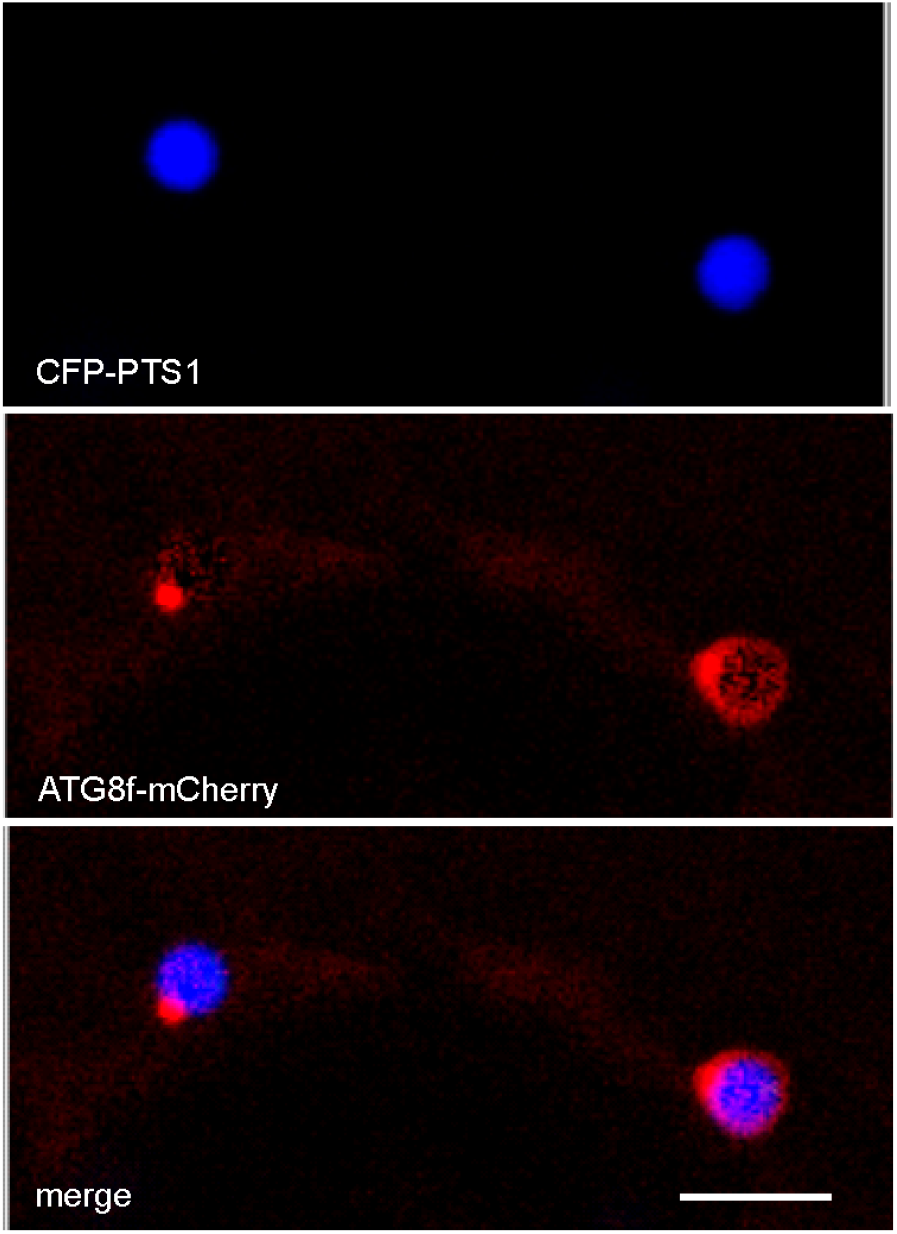
Co-expression of markers of peroxisomes (CFP-PTS1) and autophagosomes (ATG8f-mCherry). This picture shows a rarely observed advanced pexophagy with the whole peroxisome engulfed by the autophagosomal membrane (right peroxisome) and a more frequently observed association of small autophagosome at the peroxisomal periphery (left peroxisome). Scale bar = 5 µm.

**figure S6.**
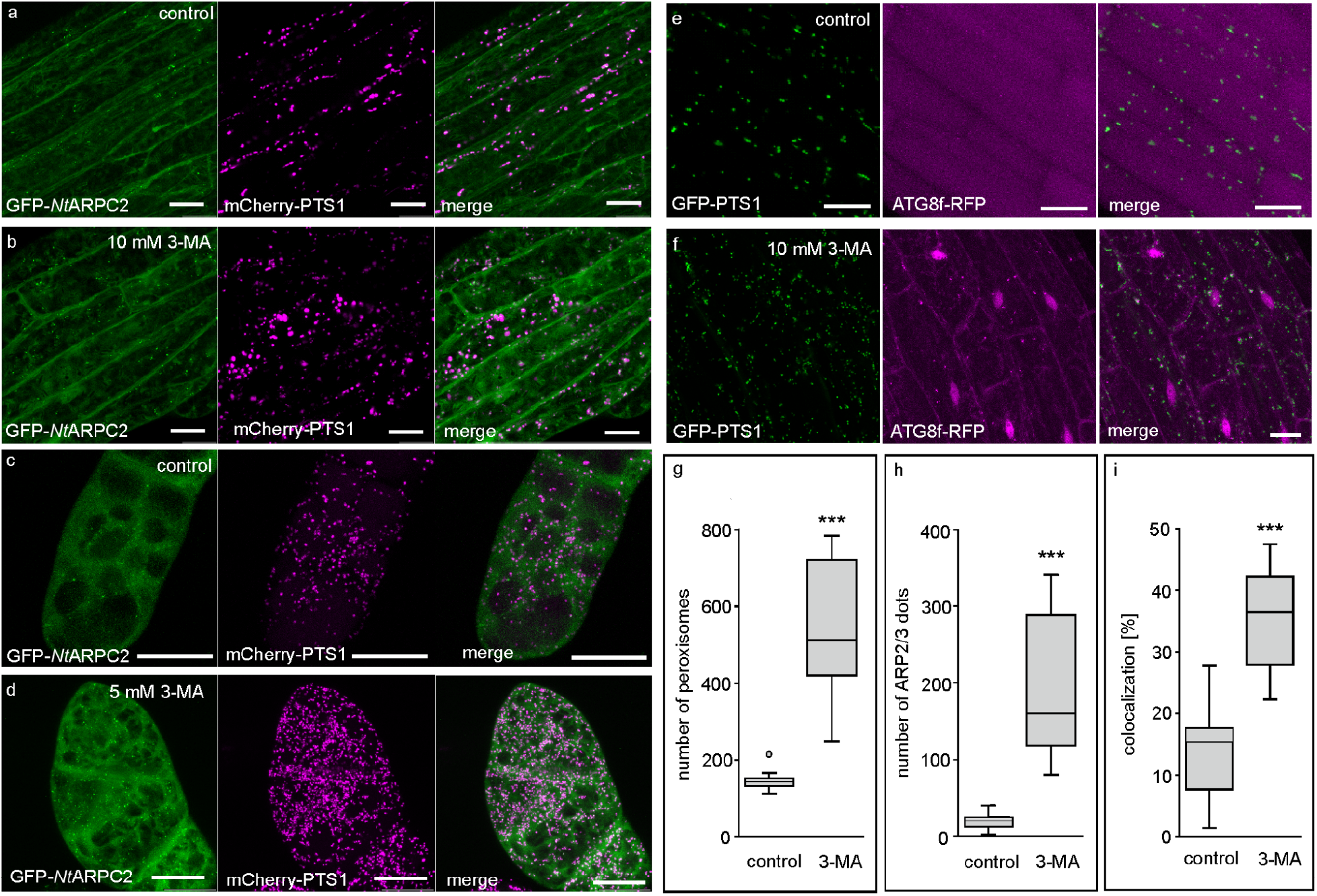
Treatment with 3-MA resulted in an increased number of both ARP2/3 dots (**a-d**) and peroxisomes (**a-f**) in both *Arabidopsis* (**a, b, e, f)** and BY-2 cells (**c, d**). (**e, f**) 3-MA treatment resulted in an increased number of autophagosomes and a change in the marker distribution. Under control conditions, the ATG8f signal was localized mainly in the vacuole, and some ATG8f-labeled structures were found in the cytoplasm (**e**). After 3-MA treatment, the ATG8f marker was found mainly in the cytoplasm and nuclei (**f**). (**g-i**) Quantification showed a significant increase in the number of peroxisomes (**g**) and ARP2/3 dots (**h**) in 3-MA treated BY-2 cells. The ratio of ARP2/3 dots to peroxisomes shows a significant increase of peroxisomes associated with ARP2/3 dots after 3-MA treatment (**i**). Quantification was performed from z-stacks of confocal pictures of fixed BY-2 cells. Central lines in boxplots represent median, n = 12 (control) and 13 (3-MA). The significance of T-tests is indicated by asterisks (*p* < 0.001 =***) Scale bars = 20 µm.

