## Supporting Information 1 for "ARP2/3 complex associates with peroxisomes to participate in pexophagy in plants"

### Protein Digestion

IP samples were resuspended in 100mM TEAB containing 2% SDC. Cysteines
were reduced with 5mM final concentration of TCEP (60 °C for 60 min) and blocked with 10mM final concentration of MMTS (10 min Room Temperature). Samples were cleaved on beads with 1 µg of trypsin at 37 °C overnight. After digestion, samples were centrifuged and supernatants were collected and acidified with TFA to 1% final concentration. SDC was removed by extraction to ethyl acetate (Masuda *et al.*, 2008). Peptides were desalted on the Michrom C18 column.

### nLC-$MS^2$ Analysis

Nano Reversed phase column (EASY-Spray column, 50 cm x 75 µm ID, PepMap C18, 2 µm particles, 100 Å pore size) was used for LC/MS analysis. Mobile phase buffer A was consisting of water and 0.1% formic acid. Mobile phase B was consisting of acetonitrile and 0.1% formic acid. Samples were loaded onto the trap column (Acclaim PepMap300, C18, 5 µm, 300 Å Wide Pore, 300 µm x 5 mm, 5 Cartridges) for 4 min at 15 µl/min. Loading buffer was consisting of water, 2% acetonitrile and 0.1% trifluoroacetic acid. Peptides were eluted with Mobile phase B gradient from 4% to 35% B in 60 min. Eluting peptide cations were converted to gas-phase ions by electrospray ionization and analyzed on a Thermo Orbitrap Fusion (Q-OT-qIT, Thermo). Survey scans of peptide precursors from 400 to 1600 m/z were performed at 120K resolution (at 200 m/z) with a 5 × 10 5 ion count target. Tandem MS was performed by isolation at 1.5 Th with the quadrupole, HCD fragmentation with a normalized collision energy of 30, and rapid scan MS analysis in the ion trap. The $MS^2$ ion count target was set to $10^4$ and the max injection time was 35 ms. Only those precursors with charge state 2–6 were sampled for $MS^2$. The dynamic exclusion duration was set to 45 s with a 10ppm tolerance around the selected precursor and its isotopes. Monoisotopic precursor selection was turned on. The instrument was run in top speed mode with 2 s cycles (Hebert *et al.*, 2014).

All data were analyzed and quantified with the MaxQuant software (version 1.5.3.8) (Cox *et al.*, 2014). The false discovery rate (FDR) was set to 1% for both proteins and peptides and we specified a minimum length of seven amino acids. The Andromeda search engine was used for the MS/MS spectra search against the *Arabidopsis thaliana* database (downloaded from Uniprot on July 2016, containing 51 445 entries). Enzyme specificity was set as C-terminal to Arg and Lys, also allowing cleavage at proline bonds and a maximum of two missed cleavages. Dithiomethylation of cysteine was selected as fixed modification and N- terminal protein acetylation and methionine oxidation as variable modifications. The “match between runs” feature of MaxQuant was used to transfer identifications to other LC-MS/MS runs based on their masses and retention time (maximum deviation 0.7 min) and this was also used in quantification experiments. Quantifications were performed with the label-free algorithms described recently. Data analysis was performed using Perseus 1.5.2.4 software
