## Supporting Table S2 for "ARP2/3 complex associates with peroxisomes to participate in pexophagy in plants"

Supporting Table S2
Primers used for cloning described constructs. Restriction sites overghangs in red.

| gene | direction | restriction site | Sequence |
|---|---|---|---|
| ARPC2 | Forward | *BamHI* | GGATCCAATGATACTATTGCAGTCACATTC |
| | Reverse | *HindIII* | AAGCTTCTACTTCGAGTTGGTGTGATTG |
| ARPC5 (GFP) | Forward | *BamHI* | GGATCCAATGGCAGAATTCGTTGAAGCTG |
| | Reverse | *HindIII* | AAGCTTTCAAACGGTGTTGATGGTATCAGTAAG |
| ARPC5 (mCh) | Forward | *BamHI* | AAAGGATCCATGGCAGAATTCGTTGAAG |
| | Reverse | *XmaI* | TTTCCCGGGAACGGTGTTGATGGTATCAG |
| MCD | Forward | *BamHI* | AAAAAAGGATCCATGAGCAAGAAAAATCTAGCGATTC |
| | Reverse | *EcoRI* | AAAAAAGAATTCCTAGAGACGGGAATGGATAC |
