## Supporting Table S1 for "ARP2/3 complex associates with peroxisomes to participate in pexophagy in plants"

Consensus protein interactors of GFP-*Nt*ARPC2, identified in both repetitions of proteomic analysis.

| Protein ID (Uniprot) | FASTA headers | PTS | intensity 1 (max. 26.3574) | intensity 2 (max. 26.1078) |
|---|---|---|---|---|
| **A0A1S4DR85**<br>**A0A1S3ZA87** | Actin-related protein 2/3 complex subunit OS<br>Actin-related protein 2/3 complex subunit OS | | 21.9136 | 23.7485 |
| **G0WLG3** | Arp2/3 complex 34 kDa subunit OS | | 26.3574 | 26.1078 |
| **A0A1S3Z3H1**<br>**A0A1S3WXP2** | Actin-related protein 2/3 complex subunit 3 OS<br>Actin-related protein 2/3 complex subunit 3 OS | | 21.5897 | 21.9216 |
| **A0A1S4BTZ8** | Actin-related protein 2/3 complex subunit 4 OS | | 21.176 | 24.4566 |
| **A0A1S3ZCR5** | Actin-related protein 2/3 complex subunit 5 OS | | 21.4188 | 24.1816 |
| **A0A1S4DI60** | actin-related protein 3 OS | | 21.5047 | 23.0667 |
| **A0A1S4A550**<br>**A0A1S4A514**<br>**A0A1S4A4Y5**<br>**A0A1S4A581** | actin-related protein 3 isoform X4 OS<br>actin-related protein 3 isoform X3 OS<br>actin-related protein 3 isoform X2 OS<br>actin-related protein 3 isoform X1 OS | | 19.7576 | 21.8903 |
| **A0A1S4A6H4** | peroxisomal fatty acid beta-oxidation multifunctional protein AIM1-like OS | PTS1 | 22.3573 | 22.0314 |
| **A0A1S4DNE8** | peroxisomal fatty acid beta-oxidation multifunctional protein AIM1-like OS | PTS1 | 20.9527 | 20.1321 |
| **A0A1S3XYS3** | peroxisomal fatty acid beta-oxidation multifunctional protein AIM1-like OS | PTS1 | 19.5933 | 22.0314 |
| **A0A1S4D439** | LOW-QUALITY PROTEIN: glyoxysomal fatty acid beta-oxidation multifunctional protein MFP-a-like OS | PTS1 | 20.2187 | 19.6894 |
| **A0A1S4D6S9**<br>**A0A1S4D762**<br>**A0A1S4D720**<br>**A0A1S4A2D2** | 4-coumarate--CoA ligase-like 6 isoform X3 OS<br>4-coumarate--CoA ligase-like 6 isoform X2 OS<br>4-coumarate--CoA ligase-like 6 isoform X1 OS<br>4-coumarate--CoA ligase-like 6 OS | PTS1 | 21.3696 | 19.7423 |
| **A0A1S4AJS0** | 4-coumarate--CoA ligase-like 7 OS | PTS1 | 24.8577 | 22.5521 |
| **A0A1S4DHZ4**<br>**A0A1S4DQN6** | probable quinone oxidoreductase OS<br>probable quinone oxidoreductase OS | PTS1 | 24.2513 | 21.5504 |
| **A0A1S3Y9E9**<br>**A0A1S4D3Y8**<br>**A0A1S4ASM7**<br>**A0A1S3YQG6** | Aspartate aminotransferase OS<br>Aspartate aminotransferase OS<br>Aspartate aminotransferase OS<br>Aspartate aminotransferase OS | PTS2 | 23.703 | 20.0947 |
| **A0A1S3Z7C4**<br>**A0A1S3Z7V8** | delta(3,5)-Delta(2,4)-dienoyl-CoA isomerase, peroxisomal isoform X1 OS<br>delta(3,5)-Delta(2,4)-dienoyl-CoA isomerase, peroxisomal isoform X2 OS | PTS1 | 23.4051 | 21.0945 |
| **A0A1S3Y1Q5**<br>**A0A1S3YTH0** | acyl-coenzyme A thioesterase 13-like OS<br>acyl-coenzyme A thioesterase 13-like OS | PTS1 | 21.8947 | 18.5317 |
| **A0A1S3YP09** | glyoxysomal processing protease, glyoxysomal-like OS | PTS1 | 24.5543 | 21.6784 |
| **A0A1S4BMM1**<br>**A0A1S3XIF9** | gamma-glutamyl peptidase 5-like OS<br>gamma-glutamyl peptidase 5-like OS | | 22.0176 | 19.499 |
